# Innate immune receptor C5aR1 regulates cancer cell fate and can be targeted to improve radiotherapy in tumours with immunosuppressive microenvironments

**DOI:** 10.1101/2023.01.10.521547

**Authors:** Callum Beach, David MacLean, Dominika Majorova, Stavros Melemenidis, Dhanya K. Nambiar, Ryan K. Kim, Gabriel N Valbuena, Silvia Guglietta, Carsten Krieg, Damavandi Mahnaz Darvish, Tatsuya Suwa, Alistair Easton, Enric Domingo, Eui Jung Moon, Dadi Jiang, Yanyan Jiang, Albert C Koong, Trent M. Woodruff, Edward E. Graves, Tim Maughan, Simon J. A. Buczacki, Manuel Stucki, Quynh-Thu Le, Simon J. Leedham, Amato J. Giaccia, Monica M Olcina

**Author notes:** Joint author (Callum Beach, David MacLean, Dominika Majorova). Corresponding Author: Monica M. Olcina, Current address: Oxford Institute of Radiation, Oncology University of Oxford, Old Road Campus Research Building, Roosevelt Drive, Oxford, OX37DQ.

## Abstract

An immunosuppressive microenvironment causes poor tumour T-cell infiltration and is associated with reduced patient overall survival in colorectal cancer. How to improve treatment responses in these tumours is still a challenge. Using an integrated screening approach to identify cancer-specific vulnerabilities, we identify complement receptor C5aR1 as a druggable target which when inhibited improves radiotherapy even in tumours displaying immunosuppressive features and poor CD8+ T-cell infiltration. While C5aR1 is well-known for its role in the immune compartment, we find that C5aR1 is also robustly expressed on malignant epithelial cells, highlighting potential tumour-cell specific functions. C5aR1 targeting results in increased NF-κB-dependent apoptosis specifically in tumours and not normal tissues; indicating that in malignant cells, C5aR1 primarily regulates cell fate. Collectively, these data reveal that increased complement gene expression is part of the stress response mounted by irradiated tumours and that targeting C5aR1 can improve radiotherapy even in tumours displaying immunosuppressive features.

## Introduction

The composition of the tumour microenvironment (TME) impacts treatment responses in cancer (Bindea et al., 2013; Llosa et al., 2015; Tauriello et al., 2018). Immunosuppressive TME features, which act as a barrier to extensive CD8+ T cell infiltration typically characterise immune cold tumours which are associated with poor prognosis (Galon et al., 2006; Guinney et al., 2015). Indeed, low density of total T lymphocytes (CD3+) at the centre or invasive margins of tumours is associated with reduced overall survival in colorectal cancer (CRC)(Galon et al., 2006). Improving treatment responses for those patients with poor tumour lymphocyte infiltration remains a challenge.

Approximately one third of all colorectal cancers arise in the rectum. Locally advanced rectal cancers are typically treated with neoadjuvant chemoradiotherapy (nCRT) prior to surgery. Unfortunately, despite nCRT leading to complete pathological regression in 20-30% of these patients, 70-80% will fail to achieve complete responses. There is therefore a need to further improve treatment responses in a significant portion of patients receiving nCRT (Cercek et al., 2018; Li et al., 2022; van der Sluis et al., 2019). Identifying targets that modulate radiosensitivity, particularly in tumours displaying immunosuppressive features could improve treatment outcomes for the most difficult to treat tumours.

High expression of complement system components is part of the inflammatory environment of colon and rectal tumours displaying the worst survival outcomes (Becht et al., 2016; Domingo et al., 2021; Guinney et al., 2015; Krieg et al., 2022). The complement system is an ancient component of innate immunity and both canonical and non-canonical functions are increasingly being recognised as important for infection control, autoimmunity and cancer (Daugan et al., 2021a; Gros et al., 2008; Pio et al., 2013; Ricklin et al., 2010; Roumenina et al., 2019). Whether in the context of cancer treatment, complement proteins are expressed and function independently of their role in the inflammatory environment remains to be fully understood.

In this study we find that in murine models which recapitulate an immunosuppressive TME, the complement system is the first immune response pathway to be upregulated at early timepoints following irradiation. Through an integrated screening approach, we identify complement receptor C5aR1 as a druggable target, which inhibits radiation-induced cell death/apoptosis through regulation of tumour cell fate. Interestingly, these effects are not observed in untransformed intestinal organoids or normal intestinal tissues *in vivo*. Consequently, targeting C5aR1 with a clinical grade and orally active C5aR1 antagonist, PMX205, results in improved tumour radiation responses *in vivo*. Importantly, PMX205 improves tumour radiation response in several murine models, including those displaying high radiation-induced complement expression and immunosuppressive features associated with CD8+ T cell exclusion.

## Results

### Identification of radiation-responsive targets in immunosuppressive tumours

When grown subcutaneously, tumour organoids originally derived from *villin*CreER *Apc*^fl/fl^ *Kras*^G12D/+^ *Trp53*fl/fl *TrgfbrI*^fl/fl^ (AKPT) mice display TME features resembling those of CRCs associated with poor outcome **(Figures 1A-C)**. These features include stromal-rich regions with high numbers of fibroblasts and macrophages but relatively few CD8^+^ T-cells **(Figures 1A-C)**. Interestingly, we found that although irradiation was able to enhance infiltration of Tregs, macrophages, neutrophils, CD4^+^ and CD8^+^ T-cells into these tumours, such infiltration was still limited to stromal regions and did not increase intra-epithelial immune cells **(Figures 1A-B and Supplementary Figure 1A and B)**. To identify radiation-responsive pathways in the immunosuppressive microenvironment of these tumours, we performed RNA-seq analysis. Network analysis of differentially expressed pathways following irradiation (RT) indicated that the complement cascade was significantly upregulated and in fact was the top ranked pathway annotated as an “Immune System pathway” in Reactome at early timepoints post RT **(Figures 1D and E and Supplementary Figure 1C and Supplementary Table 1)**. Examples of members across all the main complement functional categories (components, pattern recognition molecules, proteases, receptors and regulators) were induced following RT, with most genes showing transient enhanced expression (**Figure 1F-J**). Overall, these data suggest that complement gene expression is globally, although transiently, induced following irradiation in this model of an immunosuppressive TME.

**Figure 1:**
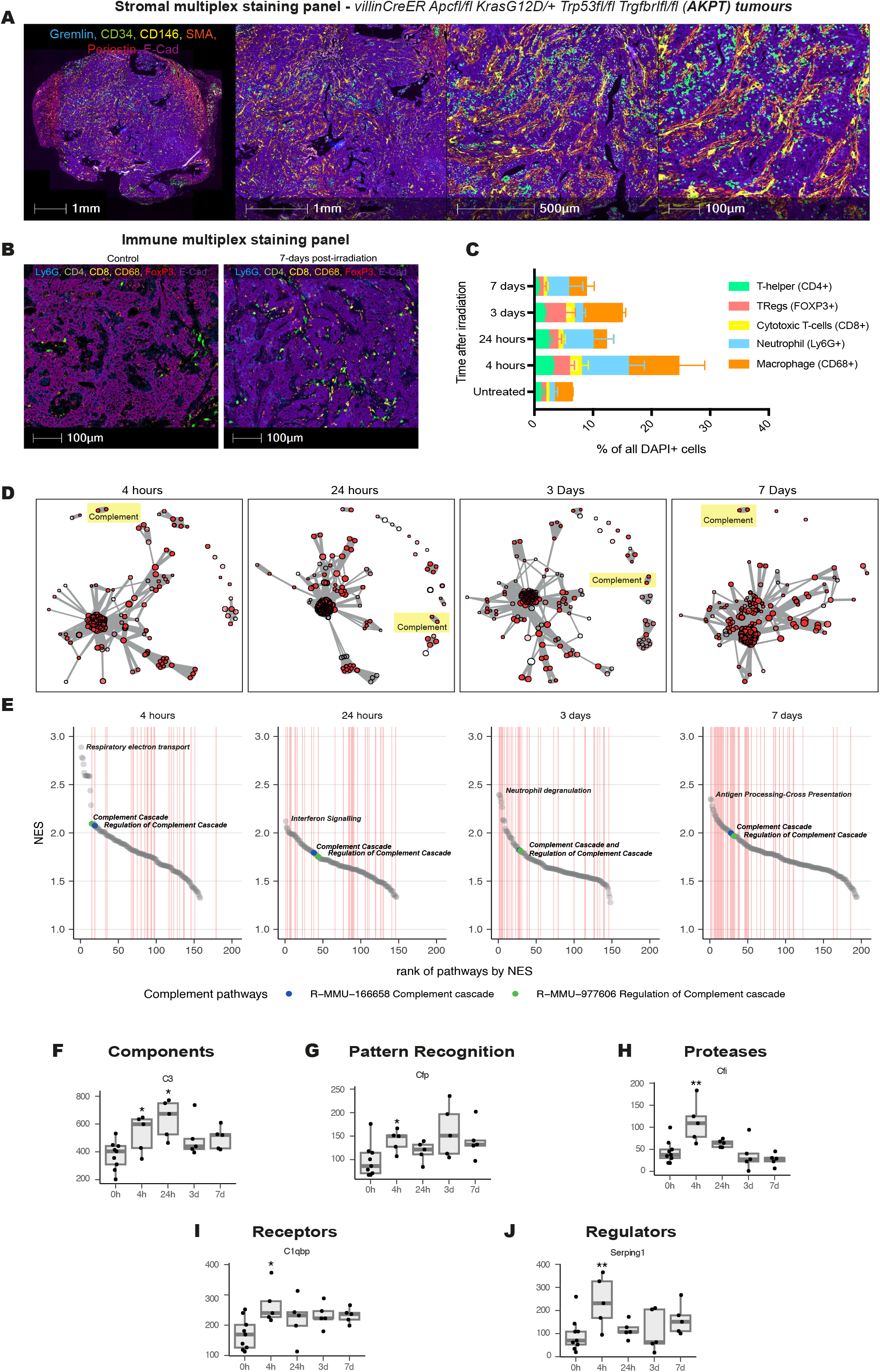
Identification of radiation-responsive targets in immunosuppressive tumours. **(A)** Representative image of *villin*CreER *Apc*^fl/fl^ *Kras*^G12D/+^ *Trp53*fl/fl *TrgfbrI*^fl/fl^ (AKPT) colorectal tumour organoids grown subcutaneously. Multiplex staining of epithelial and stromal cells is shown in whole tumour (left) and zoomed in regions. **(B)** Representative images of *villin*CreER *Apc*^fl/fl^ *Kras*^G12D/+^ *Trp53*fl/fl *TrgfbrI*^fl/fl^ (AKPT) colorectal tumour organoids grown subcutaneously and treated with either 0 Gy (left) or 15 Gy (right). Multiplex staining of epithelial and immune cells is shown. **(C)** Machine learning-based quantification of immune cell infiltration following multiplex staining at different timepoints following irradiation. **(D)** Network maps of differentially expressed pathways following RNA-sequencing (RNA-seq) of AKPT tumours harvested at different timepoints following irradiation. Significantly-enriched Reactome pathways for each timepoint are visualised as network graphs using the Fruchterman-Reingold algorithm. Each node is a pathway gene set, with the size of the nodes being proportional to the size of the gene set and pathways with lower *PFDR* appearing as less transparent nodes. Connections between nodes depend on the proportion of overlapping genes between two gene sets. The Reactome pathways related to Complement are highlighted in yellow. **(E)** The ranked Normalised Enrichment Scores (NES) are shown below the network graphs for the significant positively-enriched pathways. The complement pathways appear among the top most enriched pathways at each timepoint after 15 Gy radiation treatment. Ranks of pathways annotated as Immune System pathways in Reactome are denoted by the vertical lines in red. **(F)–(J)** Plots show expression of representative differentially expressed complement genes at different timepoints following RNA-seq of AKPT tumour receiving 15 Gy irradiation.

### C5aR1 is a radiation-responsive druggable target

To identify potential targets within the complement cascade that could be therapeutically inhibited, we queried the CanSAR database (querying complement system components, receptors, proteases and regulators). A gene was only considered a hit if it was “druggable” based on structural and ligand-based assessment as shown in green **(Figure 2A and Supplementary Table 2)**. We also interrogated the DepMap database (which combines data from CRISPR-Cas9 and RNAi screens in more than 700 cell lines), to identify cancer-derived complement proteins that may have autocrine functions specifically impacting cell fate under stress conditions. We reasoned that looking for non-essential hits would allow the identification of genes providing stress-specific dependencies and, therefore, potential therapeutic targets less likely to mediate toxicity in normal tissues. Following the combined CanSAR and Depmap analysis, we found three hits: *C5, C5AR1, C4BPA* **(Figure 2A and Supplementary Table 2)**. *ATR* was included in the screen as a positive control for essential genes since its deletion is lethal in several cell lines due to its role in replication and the DNA damage response (Cimprich and Cortez, 2008; Flynn and Zou, 2011). Interestingly, *C1QBP,* a complement gene which was recently shown to play a role in the DNA damage response by modulating DNA resection, was essential in a number of cell lines, further validating our screening approach (Bai et al., 2019) **(Figure 2A)(Supplementary Table 2)**.

**Figure 2:**
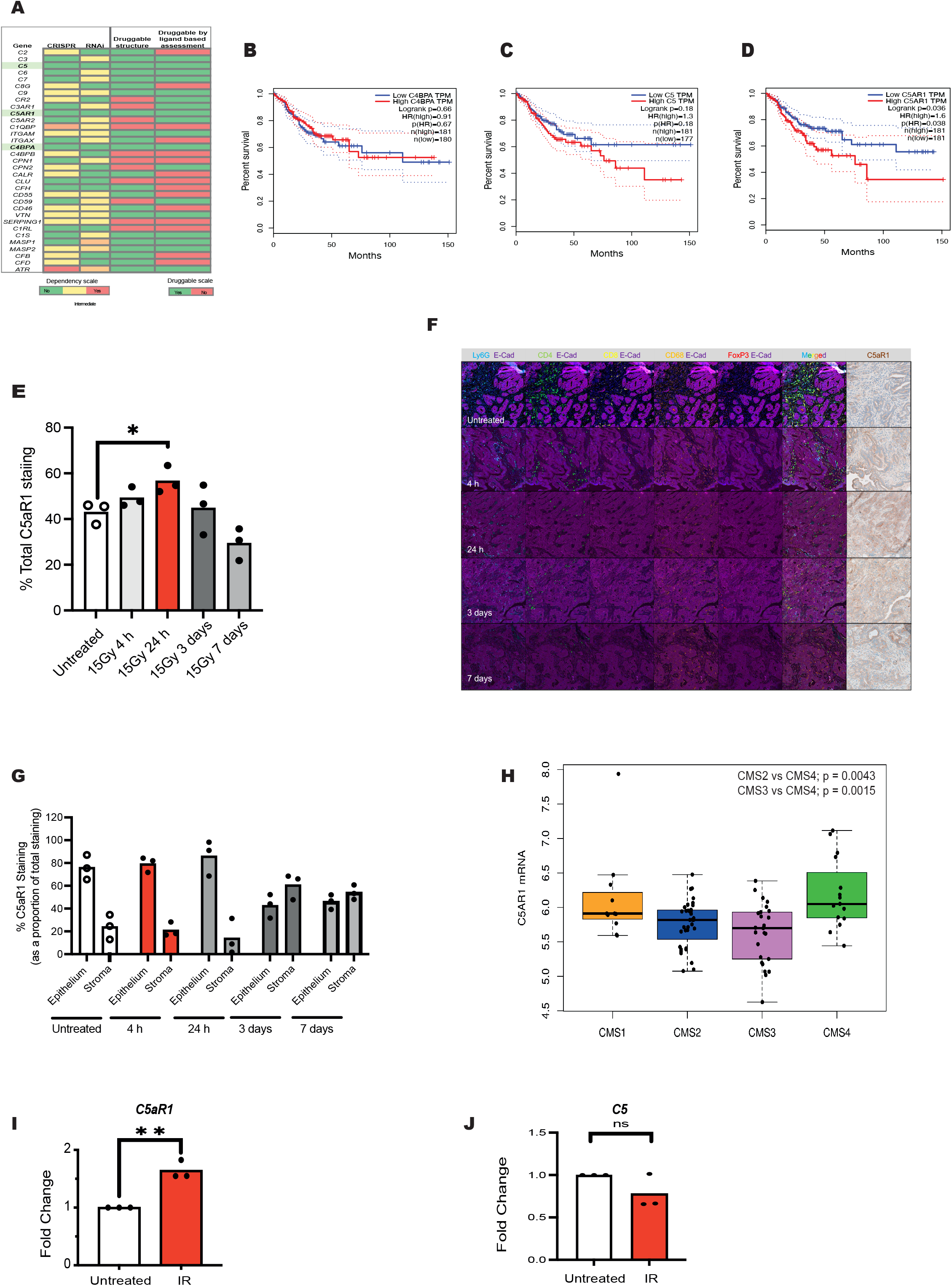
C5aR1 is a radiation-responsive druggable target. **(A)** *In silico* screen of complement genes. Data acquired from DepMap portal (https://depmap.org) Essential genes are shown in red. Non-essential genes across all studies are shown in green. Yellow/Orange = intermediate dependence or essentiality. For target tractability: green corresponds to druggable structure = Yes and druggable by ligand-based assessment = Yes. Red corresponds to druggable structure = No and druggable by ligand-based assessment = No. **(B)** GEPIA Kaplan-Meier (KM) curve for disease-free survival of TCGA colorectal cancer patients with high (red) or low (blue) C4BPA mRNA expression levels is shown. Group Cutoff = Median (http://gepia.cancer-pku.cn). **(C)** GEPIA Kaplan-Meier (KM) curve for disease-free survival of TCGA colorectal cancer patients with high (red) or low (blue) C5 mRNA expression levels is shown. Group Cutoff = Median (http://gepia.cancer-pku.cn). **(D)** GEPIA Kaplan-Meier (KM) curve for disease-free survival of TCGA colorectal cancer patients with high (red) or low (blue) C5AR1 mRNA expression levels is shown. Group Cutoff = Median (http://gepia.cancer-pku.cn). **(E)** Quantification of C5aR1 immunohistochemistry staining in AKPT tumours harvested at the indicated timepoints following irradiation. *=p<0.05. Points indicate individual mice per group. **(F)** Representative images of multiplex and C5aR1 IHC staining in untreated (unirradiated) AKPT tumours and tumours harvested 4 h, 24 h, 3 days and 7 days after RT. The different markers assessed by multiplex IHC are shown above the panels. In all fluorescent images DAPI is shown in dark blue. **(G)** Machine learning-based quantification of % of C5aR1 staining in the epithelium and stroma of AKPT tumours harvested at the indicated timepoints following irradiation. Regions of the tumour were categorised as epithelium or stroma by the HALO software using analysis of multiplex immune and E-cadherin staining as shown in (F). **(H)** C5AR1 mRNA expression in pre-treatment rectal tumour biopsies (N=129) classified according to consensus molecular subtypes (CMS). Significance assessed by Wilcoxon test. **(I)** mRNA expression of *C5AR1/housekeeping* in HCT 116 cells treated with either 0 or 9 Gy irradiation. Individual points indicate average from biologically independent replicates. n=3. **(J)** mRNA expression of *C5/housekeeping* in HCT 116 cells treated with either 0 or 9 Gy irradiation. Individual points indicate average from biologically independent replicates. n=3.

*C5* encodes a complement component, which when cleaved will form C5a. C5aR1 is the main signalling receptor for the C5a ligand. *C4BPA* encodes the alpha chain of complement regulator C4BP (Hofmeyer et al., 2013) and there are currently no known pharmacological approaches for targeting C4BPA. To further narrow down which hit would be the best therapeutic target, we assessed the association of C5, C5aR1 and C4BPA mRNA expression with prognostic outcomes and found that only high C5aR1 mRNA expression was associated with significantly poor disease-free survival in colorectal cancer **(Figure 2B-D)**. We confirmed that high C5aR1 mRNA expression was correlated with decreased overall survival in a further independent dataset **(Supplementary Figure 2A)**.

*In vivo* we found that C5aR1 was robustly expressed in AKPT tumours at baseline. A transient induction in C5aR1 expression was also observed after RT **(Figure 2E and Supplementary Figure 2B)**. We next used multiplex staining and machine learning-based image analysis software to investigate the cellular compartments expressing C5aR1 in greater detail (**Figure 2F, G and Supplementary Figure 2C**, for negative control C5aR1 staining). Interestingly, we found that at earlier timepoints post RT C5aR1 expression is more prominent in the epithelium, while stromal C5aR1 expression appears to dominate at later timepoints (3 and 7 days)(**Figure 2G**). The dominance of stromal C5aR1 expression at these later timepoints may reflect increased infiltration of C5aR1-expressing immune cells following irradiation, although this remains to be formally assessed. AKPT tumours harbour stromal infiltration features comparable to those patients with tumours classified as consensus molecular subtype (CMS) 4. We therefore asked whether C5aR1 expression might be differentially expressed across subtypes (CMS1-4). Analysis of pre-treatment rectal tumour biopsies identified that those classified as CMS4 had the highest RNA levels of C5aR1 expression compared to the other subtypes **(Figure 2H**).

To directly assess whether radiation could impact tumour cell intrinsic expression of C5aR1 or C5, we turned to an *in vitro* system. Following treatment of mouse and human colorectal tumour cells with irradiation, we found a modest but reproducible increase in C5aR1 (but not C5) across cell lines **(Figure 2I and J and Supplementary Figures 2E-G)**.

### C5aR1 regulates tumour cell survival under stress

PMX205 is a selective inhibitor of C5aR1 currently undergoing clinical testing for ALS where it is reportedly well-tolerated (Kumar et al., 2020). We assessed the effects of treating colorectal tumour cells with PMX205. As anticipated, given the fact that C5aR1 is a G-protein coupled receptor (GPCR), we found reduced ERK1/2 and RelA phosphorylation (as a readout for NFκB signalling) in PMX205 treated cells **(Figure 3A)**. However, PMX205 had negligible effects on AKT phosphorylation (at Threonine 308) **(Supplementary Figure 3A and B)**.

**Figure 3:**
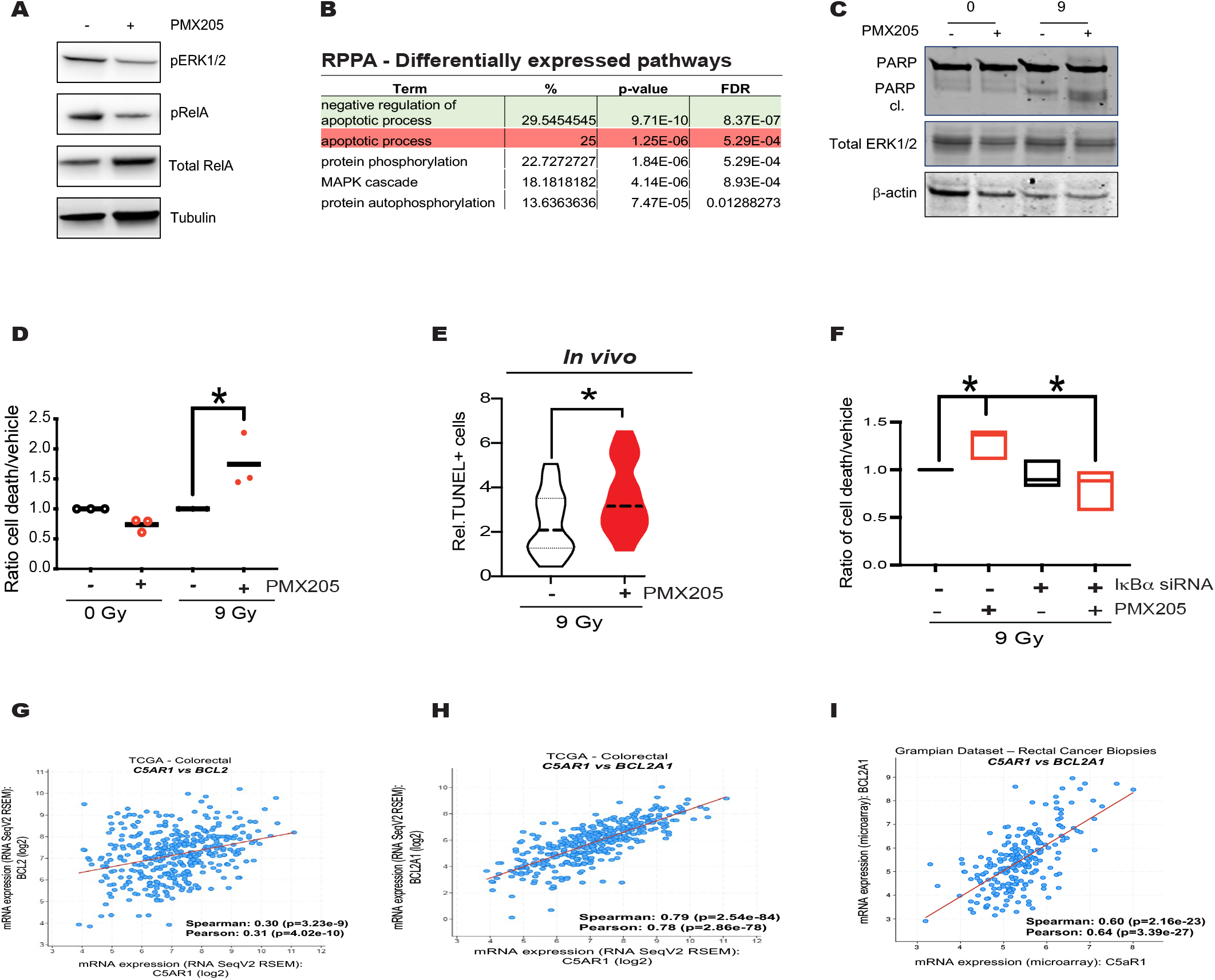
C5aR1 regulates tumour cell survival under stress. **(A)** HCT116 cells were treated with either vehicle or PMX205 for 48 hours. Western blotting was carried with the antibodies indicated. Tubulin was used as the loading control. n=3 **(B)** Table of top differentially expressed pathways from the RPPA shown in (A). Apoptotic processes shaded in green include proteins that are downregulated, those in red include upregulated proteins. The % of enriched terms, p-value and false discovery rate (FDR) are shown. **(C)** HCT116 cells were treated with 0 or 9 Gy and either vehicle or PMX205 1 hour before irradiation. Cells were harvested 48 hours post-IR. Western blotting was carried with the antibodies indicated. β-actin was used as the loading control. **(D)** The graph represents the ratio of dead/apoptotic cells expressed as a % of the whole population relative to vehicle treated cells for the experiment shown in (C). n=3. * = p<0.05, two-tailed t-test. **(E)** Graph shows TUNEL+ cells per field of view of sections from HCT116 subcutaneous tumours grown in athymic nude mice. Mice were treated with 9 Gy single dose irradiation and either vehicle or PMX205 treatment for 3 doses flanking the irradiation dose (on day 0, 1 and 2). * = p<0.05 by two tailed t-test. **(F)** The graph represents the number of dead/apoptotic cells expressed as a % of the whole population for HCT 116 cells transfected with either Scr or IκBα siRNA and treated with either vehicle or PMX205 for 1 hour before irradiation with either 0 or 9 Gy IR. Cells were harvested 48 hours post-IR. Independent fields of view from a representative experiment are show, n=3. * = p<0.05 by two tailed t-test. **(G)** Correlation of *BCL2* and *C5AR1* mRNA expression in TCGA colorectal patient samples. Spearman and Pearson correlations are show. Data accessed through cBioportal.org. **(H)** Correlation of *BCL2A1* and *C5AR1* mRNA expression in TCGA colorectal patient samples. Spearman and Pearson correlations are show. Data accessed through cBioportal.org. **(I)** Correlation of *BCL2A1* and *C5AR1* mRNA expression in Grampian rectal cancer biopsies Spearman and Pearson correlations are show. Data accessed through SCORT cBioportal.org.

Changes in MAPK signalling could impact cell cycle distribution which in turn could impact cellular radiosensitivity. However, we did not note any significant differences in cell cycle profiles by flow cytometry in cells treated +/− PMX205 +/− RT **(Supplementary Figure 3C)**. We also did not note significant changes in γH2AX levels between treatment groups (at late timepoints) suggesting that DNA repair is likely unaffected by PMX205 **(Supplementary Figure 3D).** Functional annotation analysis of differentially expressed proteins following reverse-phase protein array (RPPA) analysis, indicated that the top differentially expressed pathways at the protein level clustered around apoptosis/cell death with negative regulators of apoptosis being repressed following PMX205 treatment and positive regulators of apoptosis being upregulated **(Figure 3B, Supplementary Figure 3E and Supplementary Tables 3 and 4)**. Protein phosphorylation and MAPK cascade were also differentially expressed, which would be consistent with reduced GPCR activity downstream of C5aR1 inhibition (by PMX205) and our western blotting data **(Figure 3A and B)**. In line with the RPPA data, PMX205 treatment resulted in increased apoptosis in tumour cells following irradiation **(Figures 3C, D and Supplementary Figure 3F).** C5aR1 depletion also resulted in increased apoptosis following irradiation. **(Supplementary Figure 3G and H).** Targeting C5aR1 did not result in increased apoptosis in the absence of irradiation, in agreement with a stress-specific role in modulating cell death. Supporting this, in HCT 116 xenografts, we observed increased apoptosis in PMX205 treated tumours following irradiation **(Figure 3E).**

To investigate whether apoptosis of PMX205-treated tumour cells occurred downstream of attenuated GPCR-associated signalling, we first depleted NF-κB inhibitor, IκBα as a means of interrogating the NF-κB dependence of the effects observed. If PMX205-mediated apoptosis was occurring in an NF-κB-dependent manner, IκBα depletion would be expected to result in decreased apoptosis in PMX205 treated cells. We indeed observed that, following irradiation, IκBα depletion in PMX205 treated cells attenuated the apoptotic response **(Figure 3F and Supplementary Figures 3I and J)**. IκBα depletion did not have a dramatic effect on apoptosis levels in the vehicle + RT treated cells. We hypothesize this is because IκBα levels are reduced by DNA damaging agents, and NF-κB signalling is already high in these cells. We also depleted RelA and found that while RelA depletion increased apoptosis levels in RT + vehicle treated cells (as expected); there was no further increase in apoptosis in RT + PMX205 treated cells (presumably since these cells already have reduced “active” RelA **(Supplementary Figures 3K and L)**. Furthermore, interrogation of the TCGA patient datasets where high C5aR1 mRNA expression was associated with poor outcome, identified that C5aR1 expression was positively and significantly correlated with pro-survival/anti-apoptosis genes including NF-κB target genes in the BCL2 family **(Figures 3G and H**). We confirmed these associations in two further independent patient datasets including rectal cancer patient biopsies collected prior to radiotherapy **(Figure 3I and Supplementary Figure 3M)**. Treatment of colorectal cancer cells with PMX205 also resulted in reduced mRNA expression of BCL2 **(Supplementary Figure 3N).**

We next assessed the effects of ERK inhibition on PMX205-induced apoptosis by treating colorectal cancer cells with PMX205 and ERK inhibitor Selumetinib (with or without irradiation). As expected, PMX205 and Selumetinib alone resulted in enhanced apoptosis following irradiation; with PMX205 displaying the most significant effects **(Supplementary Figure 3O and P)**. A moderate (yet not significant) increase in apoptosis was also observed when both compounds were combined **(Supplementary Figure 3O and P)**. These data suggest that although attenuated ERK may contribute to apoptosis following PMX205 treatment, it is unlikely to be a main driver of the apoptotic effect observed. Together, these data suggest that C5aR1 mediates tumour cell pro-survival signalling, with NF-κB acting as a key regulator of this response. In support of this conclusion, increased cell death following PMX205 treatment occurs downstream of attenuated NF-κB signalling.

### C5aR1 deficiency does not result in increased apoptosis in healthy intestinal epithelium

To assess whether the cell-intrinsic effects of complement were specific to malignant cells we first considered if organoids derived from different genotypes expressed complement genes when grown *in vitro* (and therefore in the absence of systemic complement or TME-derived complement products). Interestingly, following RNA-seq we found that complement genes were significantly differentially expressed in both APKT tumour organoids as well as organoids derived from villinCreER Kras^G12D/+;^Trp53^fl/fl^ Rosa26^N1icd/+^ (KPN) mice (compared to untransformed WT organoids). Over 74% of the complement genes queried were expressed at significantly higher levels in the AKPT organoids compared to the untransformed WT organoids (including C5aR1) suggesting that increased cell intrinsic complement expression is a malignant cell-associated phenomenon **(Figure 4A and B)**.

**Figure 4:**
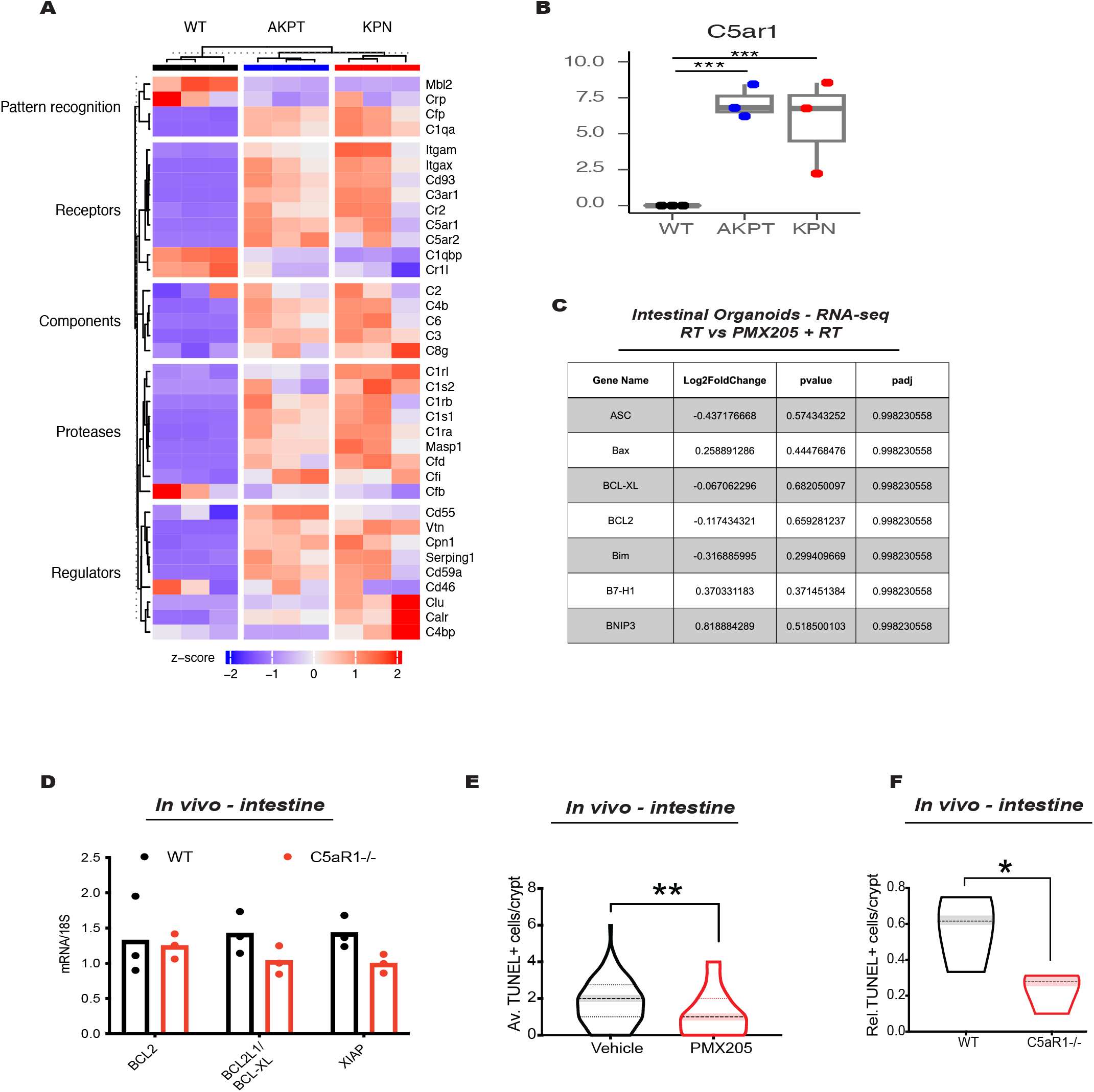
C5aR1 deficiency does not result in increased apoptosis in healthy intestinal epithelium. **(A)** Heatmap of complement genes differentially expressed by RNA-seq in WT, AKPT or KPN organoids grown *in vitro*. Upregulation is shown in red as per z-score indicated below. Data for 3 independent samples is shown from data deposited at the ArrayExpress database under accession number E-MTAB-11769 (Gil Vazquez et al., 2022). **(B)** C5aR1 expression (CPM) assessed by RNA-seq in WT, AKPT or KPN organoids grown in vitro as in (A). **(C)** Table summarising gene expression changes following RNA-seq of mouse intestinal organoids treated with either 9 Gy vs PMX205 + 9 Gy irradiation (organoids harvested 48 h following irradiation). n=3 biological replicates. **(D)** mRNA expression of *BCL2, BCL2L1 and XIAP* in WT or C5aR1^-/-^ mice treated with 9 Gy irradiation. Points indicate individual mice per group. **(E)** Graph shows the average TUNEL+ cells/crypt of mice treated with 9 Gy and either vehicle or PMX205. Intestines were harvested 72 h post irradiation. n=3 mice per group. **(F)** Graph shows the relative number of TUNEL+ cells/crypt of WT or C5aR1^-/-^ mice treated with 9 Gy. Intestines were harvested 48 h post irradiation. n=3 mice per group.

To investigate whether targeting C5aR1 might also alter pro-survival signalling in healthy tissues we assessed RelA and ERK phosphorylation in untransformed intestinal organoids. We noted that while ERK phosphorylation was reduced following PMX205 treatment, RelA phosphorylation was not affected **(Supplementary Figure 4A)**. We also performed RNA-sequencing in intestinal organoids treated +/− PMX205 +/− RT. Following gene ontology analysis, as expected, genes within the GPCR activity pathway were amongst those differentially expressed (downregulated) in organoids treated with PMX205 (**Supplementary Figure 4B**, additional differentially expressed pathways shown in **Supplementary Figure 4C)**. In line with the lack of RelA phosphorylation changes observed by western blotting, we noted that transcriptional NF-κB antiapoptosis target genes were not differentially expressed in PMX205 + RT vs RT alone treated organoids **(Figure 4C)**. Similarly, *in vivo*, small intestines did not show significant transcriptional changes in anti-apoptosis target genes in the BCL2 family with deletion of C5aR1 (following irradiation) **(Figure 4D)**. These data suggest that changes in NFκB-anti-apoptotic signalling downstream of C5aR1 do not occur in the untransformed intestinal epithelium. In line with these signalling changes, we did not observe an increase in apoptosis in small intestines following PMX205 treatment or C5aR1 loss **(Figure 4E and F)**. In fact, small intestinal crypts *in vivo* had significantly reduced apoptosis following irradiation and PMX205 treatment **(Figure 4E)**. Similarly, small intestinal crypts of C5aR1^-/-^ displayed significantly reduced apoptosis compared to WT mice following total abdominal irradiation **(Figure 4F)**. Together these data indicate that C5aR1 attenuates stress-induced apoptosis in malignant but not non-transformed epithelial cells.

### C5aR1 inhibition improves tumour radiation response

To assess whether targeting C5aR1 could improve tumour radiation responses *in vivo* we treated MC38 subcutaneous tumours with PMX205 and either no irradiation, fractionated radiotherapy (3 × 4.45 Gy) or single dose irradiation (9Gy) (as shown in three different treatment schemes in **Figure 5A**). PMX205 treatment in the absence of irradiation did not have significant effects on tumour response, consistent with a non-essential role for C5aR1 in the absence of stress **(Figure 5B)**. PMX205 treatment also had no negative effects on mouse weight **(Supplementary Figure 5A and B)**. However, following treatment with either single dose (9 Gy) or equivalent multiple fractionation (3 × 4.45 Gy) regimens (equivalent assuming an α/β ratio of 5.06) (Suwinski et al., 2007) PMX205 treatment significantly improved tumour radiation response (despite the short PMX205 dosing regimen used) **(Figures 5C and D and Supplementary Figures 5C and D)**.

**Figure 5:**
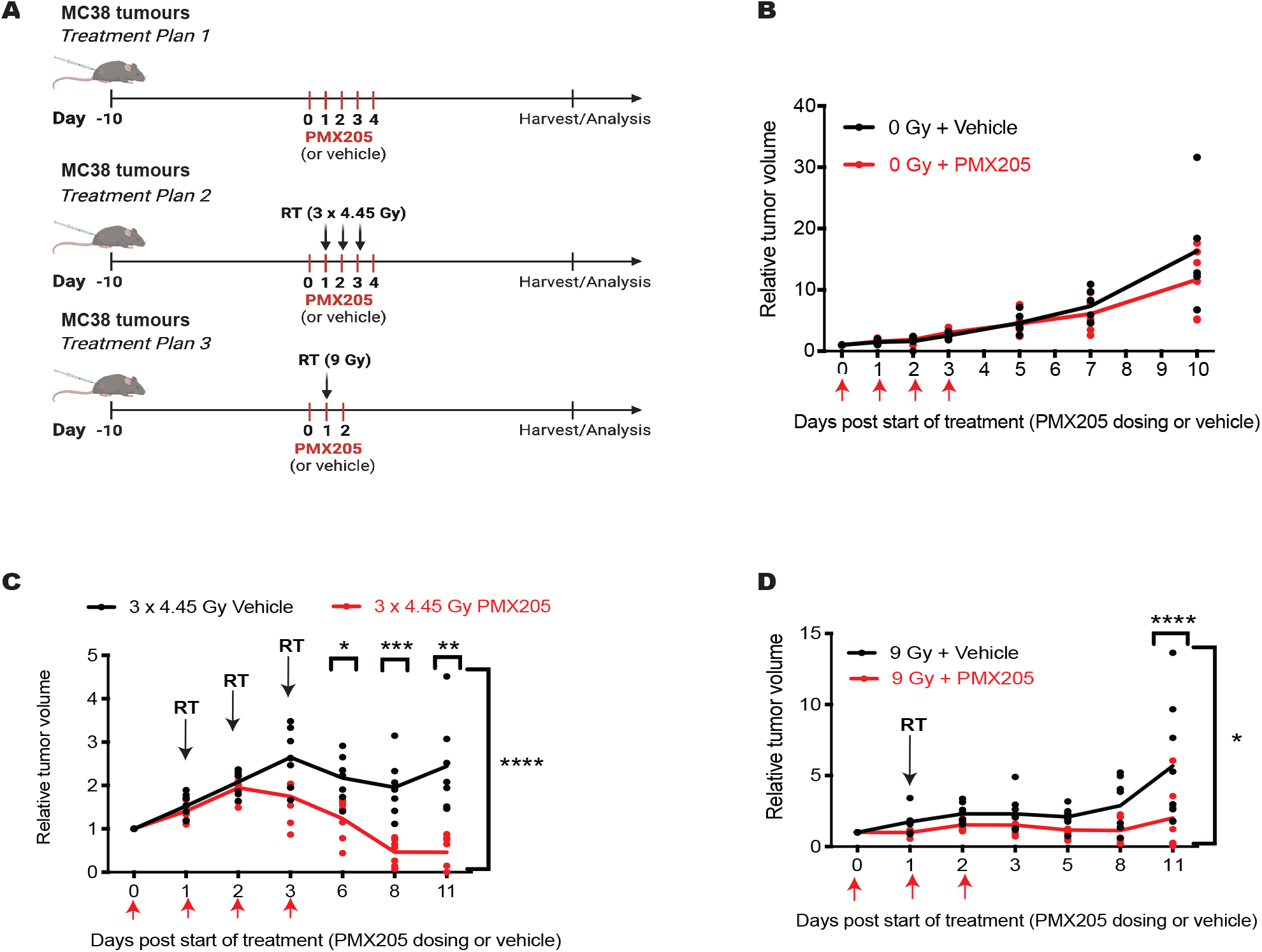
C5aR1 inhibition improves tumour radiation response. **(A)** Schematic representation of the treatment schemes followed. **(B)** Relative tumour growth curves are shown for MC38 subcutaneous tumours treated with either vehicle or PMX205 treatment for 3 doses (on day 0, 1 and 2). **(C)** Relative tumour growth curves are shown for MC38 subcutaneous tumours treated with 3 × 4.45 Gy and either vehicle or PMX205 treatment for 3 doses flanking the irradiation dose (on day 0, 1 and 2). * = p<0.05, ** = p<0.01, *** = p<0.001, comparing vehicle and PMX205 treated mice at days 6, 8 and 11 respectively. **** = p<0.0001 comparing day 0 to day 11 by 2-way ANOVA with Dunnett’s comparison test. Individual points represent individual mice per group. **(D)** Relative tumour growth curves are shown for MC38 subcutaneous tumours treated with 9 Gy single dose irradiation and either vehicle or PMX205 treatment for 3 doses flanking the irradiation dose (on day 0, 1 and 2). **** = p<0.0001 comparing vehicle and PMX205 treated mice at day 11. * = p<0.05 comparing day 0 to day 11 by 2-way ANOVA with Dunnett’s comparison test. Individual points represent individual mice per group.

### Targeting C5aR1 does not increase the % of CD8^+^ T-cells in the tumour following irradiation

Targeting C5aR1 is currently undergoing clinical testing as a means of reinvigorating anti-tumour CD8^+^ T cell responses (Massard et al., 2019). Since we had observed stromal C5aR1 expression in irradiated tumours, we next investigated tumour immune infiltration changes in tumour draining lymph nodes and tumours following PMX205 and irradiation (tissues harvested 7 days postirradiation as shown in **Figure 6A**). Interestingly, no significant changes in CD3, CD4, NK, or B-cells were found in the tumour draining lymph nodes across treatment groups **(Figures 6B-E and Supplementary Figure 6A)**. Mice treated with PMX205 alone, however, did display a higher % of CD8^+^ T cells compared to those treated with the PMX205 and irradiation combination **(Figures 6F).** Mice treated with PMX205 alone also had a reduced % of Tregs compared to vehicle treated mice **(Figure 6G)**. However, these changes did not correlate with altered functionality/effector functions of CD8^+^ T cells which showed comparable expression of IFNγ, GrzB and TNFα across all treatment groups at this timepoint **(Figures 6H-L)**. In the tumour, we found that although 9 Gy irradiation, as expected, significantly increased levels of CD3^+^ and CD8^+^ T-cells; treatment with PMX205 did not further increase the % of these cells **(Figure 6M, 6N and Supplementary Figure 6B)**. In fact, the % of CD8^+^ T-cells was significantly reduced in irradiated tumours after PMX205 treatment when compared to irradiated vehicle-treated mice **(Figure 6N)**. This was surprising given the improved tumour regression previously observed in these models **(Figure 5B)**. No significant changes in NK cells or either monocytic or granulocytic myeloid derived suppressor (M-MDSC or PMN-MDSC respectively) cells were observed across treatment groups in the tumour **(Supplementary Figure 6C-F)**. These data indicated that improved tumour radiation responses following PMX205 treatment can occur despite reduced tumour CD8^+^ T cell infiltration changes.

**Figure 6:**
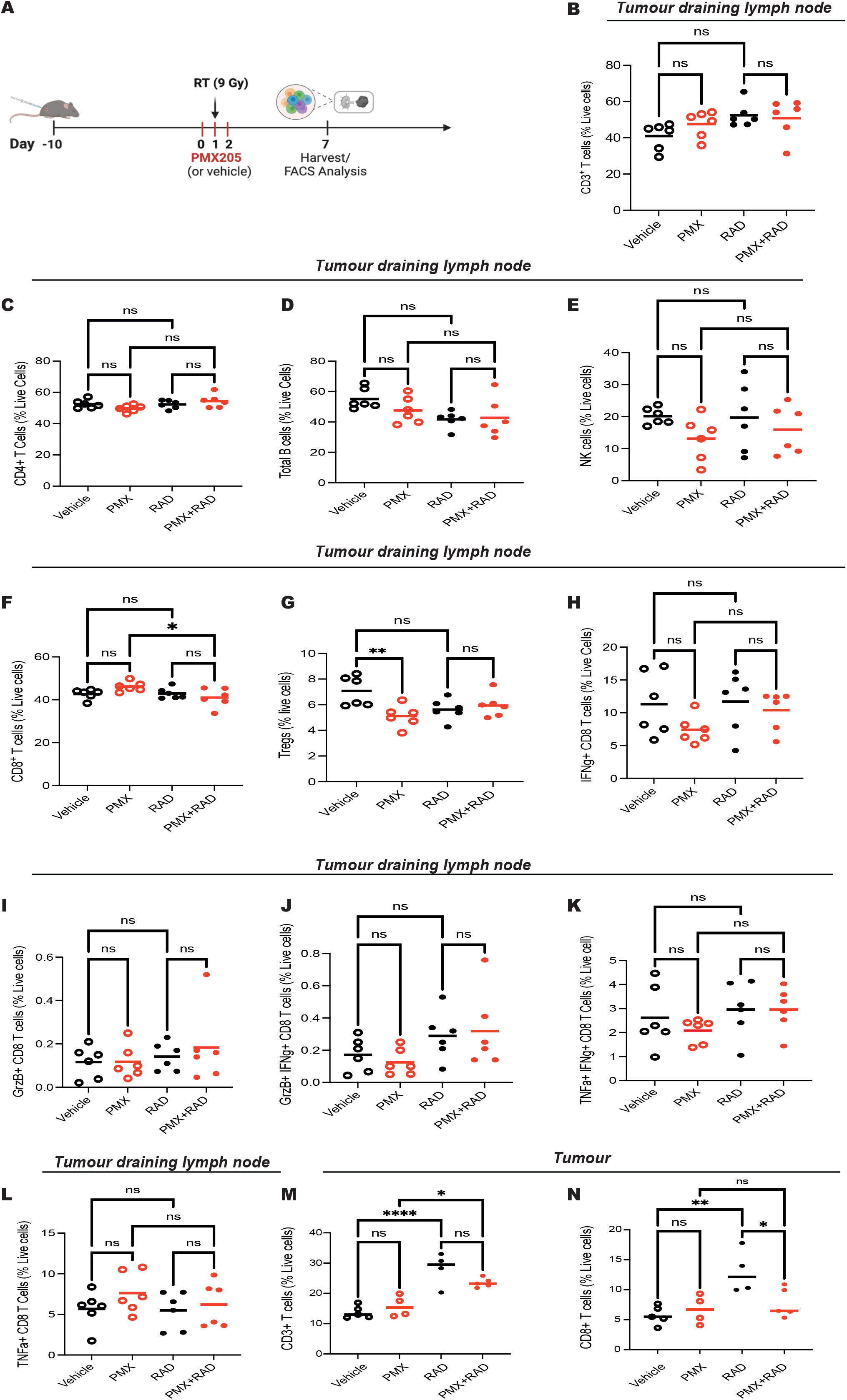
Targeting C5aR1 does not increase the % of CD8+ T-cells in the tumour following irradiation. **(A)** Schematic representation of the experimental design followed in B-N. **(B)** Graph shows CD3^+^ T-cells (as a % of live cells) in tumour draining lymph nodes of mice receiving 0 or 9 Gy and either vehicle or PMX205 treatment following the same dosing scheme as shown in A. Tumours were harvested 7 days after irradiation. ns = not significant by ordinary one-way ANOVA with Tukey’s multiple comparisons test. **(C)** Graph shows CD4^+^ T-cells (as a % of live cells) in tumour draining lymph nodes of mice receiving 0 or 9 Gy and either vehicle or PMX205 treatment following the same dosing scheme as shown in A. Tumours were harvested 7 days after irradiation. ns = not significant by ordinary one-way ANOVA with Tukey’s multiple comparisons test. **(D)** Graph shows total B cells (as a % of live cells) in tumour draining lymph nodes of mice receiving 0 or 9 Gy and either vehicle or PMX205 treatment following the same dosing scheme as shown in A. Tumours were harvested 7 days after irradiation. ns = not significant by ordinary one-way ANOVA with Tukey’s multiple comparisons test. **(E)** Graph shows total NK cells (as a % of live cells) in tumour draining lymph nodes of mice receiving 0 or 9 Gy and either vehicle or PMX205 treatment following the same dosing scheme as shown in A. Tumours were harvested 7 days after irradiation. ns = not significant by ordinary one-way ANOVA with Tukey’s multiple comparisons test. **(F)** Graph shows CD8^+^ T-cells (as a % of live cells) in tumour draining lymph nodes of mice receiving 0 or 9 Gy and either vehicle or PMX205 treatment following the same dosing scheme as shown in A. Tumours were harvested 7 days after irradiation. ns = not significant; * = p<0.05 by ordinary one-way ANOVA with Tukey’s multiple comparisons test. **(G)** Graph shows Tregs-cells (as a % of live cells) in tumour draining lymph nodes of mice receiving 0 or 9 Gy and either vehicle or PMX205 treatment following the same dosing scheme as shown in A. Tumours were harvested 7 days after irradiation. ns = not significant; ** = p<0.01 by ordinary one-way ANOVA with Tukey’s multiple comparisons test. **(H)** Graph shows IFNγ^+^ CD8 T-cells (as a % of live cells) in tumour draining lymph nodes of mice receiving 0 or 9 Gy and either vehicle or PMX205 treatment following the same dosing scheme as shown in A. Tumours were harvested 7 days after irradiation. ns = not significant by ordinary one-way ANOVA with Tukey’s multiple comparisons test. **(I)** Graph shows GrzB^+^CD8 T-cells (as a % of live cells) in tumour draining lymph nodes of mice receiving 0 or 9 Gy and either vehicle or PMX205 treatment following the same dosing scheme as shown in A. Tumours were harvested 7 days after irradiation. ns = not significant by ordinary one-way ANOVA with Tukey’s multiple comparisons test. **(J)** Graph shows GrzB^+^ IFNγ^+^ CD8 T-cells (as a % of live cells) in tumour draining lymph nodes of mice receiving 0 or 9 Gy and either vehicle or PMX205 treatment following the same dosing scheme as shown in A. Tumours were harvested 7 days after irradiation. ns = not significant by ordinary one-way ANOVA with Tukey’s multiple comparisons test. **(K)** Graph shows TNFα^+^ IFNγ^+^ CD8 T-cells (as a % of live cells) in tumour draining lymph nodes of mice receiving 0 or 9 Gy and either vehicle or PMX205 treatment following the same dosing scheme as shown in A. Tumours were harvested 7 days after irradiation. ns = not significant by ordinary one-way ANOVA with Tukey’s multiple comparisons test. **(L)** Graph shows TNFα^+^ CD8 T-cells (as a % of live cells) in tumour draining lymph nodes of mice receiving 0 or 9 Gy and either vehicle or PMX205 treatment following the same dosing scheme as shown in A. Tumours were harvested 7 days after irradiation. ns = not significant by ordinary one-way ANOVA with Tukey’s multiple comparisons test. **(M)** Graph shows CD3^+^ T-cells (as a % of live cells) in tumours of mice receiving 0 or 9 Gy and either vehicle or PMX205 treatment following the same dosing scheme as shown in A. Tumours were harvested 7 days after irradiation. ns = not significant; * = p<0.05; **** = p < 0.0001 by ordinary one-way ANOVA with Tukey’s multiple comparisons test. **(N)** Graph shows CD8^+^ T-cells (as a % of live cells) in tumours of mice receiving 0 or 9 Gy and either vehicle or PMX205 treatment following the same dosing scheme as shown in A. Tumours were harvested 7 days after irradiation. ns = not significant; * = p<0.05; ** = p < 0.01 by ordinary one-way ANOVA with Tukey’s multiple comparisons test.

### C5aR1 inhibition can improve radiotherapy in tumours with an immunosuppressive microenvironment

The improved tumour responses observed in the context of reduced tumour CD8^+^ T cell numbers, made us question whether PMX205 could be used to improve radiation responses in other models displaying low CD8^+^ T-cell infiltration. We, therefore, next asked whether PMX205 could improve tumour response in the AKPT models where we had previously observed robust C5aR1 expression and low CD8^+^ T cell tumour infiltration. In support of C5aR1 having cell intrinsic effects we observed that, *in vitro*, PMX205 reduced survival of AKPT organoids upon treatment with increasing doses of irradiation **(Figure 7A)**. When these organoids were grown as subcutaneous tumours, PMX205 alone had no significant effect on tumour growth (as previously observed in the MC38 model) **(Figure 7B and C)**. However, combination treatment with PMX205 and RT, resulted in a significant tumour growth delay, increased apoptosis and dramatic improvement in tumour-free survival with 20% of mice having unpalpable tumours at the end of the experiment **(Figure 7D-F)**. Together our data indicate that targeting C5aR1 can improve tumour radiation responses even in models with low tumour CD8^+^ T-cell infiltration. Importantly, improved responses are associated with increased tumour cell apoptosis and without concomitant increases in healthy intestinal epithelial cell apoptosis (**Figure 7G).**

**Figure 7:**
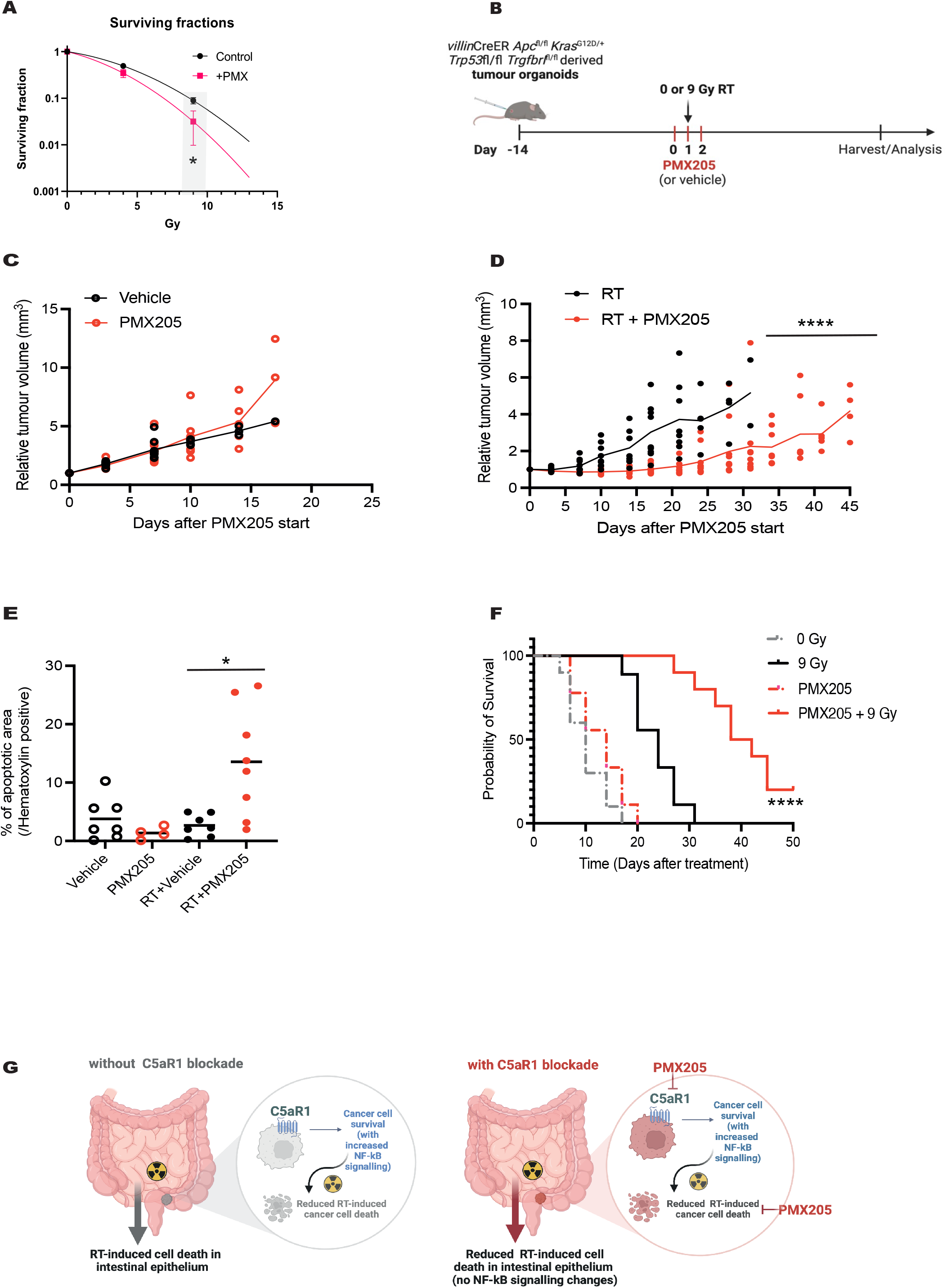
C5aR1 inhibition can improve radiotherapy in tumours with an immunosuppressive microenvironment. **(A)** Surviving fraction curve of APKT organoids treated with either vehicle or PMX205 and increasing doses of radiation. Curves represent counted viable organoids (as determined by size and structure) vs irradiation dose. **(B)** Schematic representation of the experimental design followed in C-E. **(C)** Tumour growth curves for AKPT organoids grown subcutaneously and treated with 0 Gy and either vehicle or PMX205 flanking the irradiation. Individual points = individual mice per group. **(D)** Tumour growth curves for AKPT organoids grown subcutaneously and treated with 9 Gy and either vehicle or PMX205 flanking the irradiation. Individual points = individual mice per group. **** = p<0.0001 by 2-way ANOVA with Dunnett’s comparison test. **(E)** Graph shows the percentage of apoptotic (TUNEL+) area/hematoxylin positive area in AKPT tumour bearing mice from C and D. Points indicate individual mice per group. **(F)** Survival curves of tumour-bearing mice from C and D are shown. Probability of survival as per the study endpoint criteria is shown. Curve comparison by Log-Rank **** = p<0.0001. **(G)** Schematic representation of the working model is shown. C5aR1 attenuates RT-induced cancer cell death via increased pro-survival signalling (including NF-κB). In the intestinal epithelium irradiation results in increased cell death. Upon C5aR1 blockade, tumour cells undergo increased RT-induced cell death which is not observed in the intestinal epithelium.

## Discussion

Identifying tumour-promoting components of the TME presents therapeutic opportunities. However, expression of these components can be extremely dynamic, and may be governed by selective pressures such as those posed by treatment-induced stress responses. The effects of these selective pressures on dysregulation of complement components and their evolving functions in the TME is unclear. Here we report that complement gene expression is induced after irradiation of murine tumour models which recapitulate features of human tumours displaying the worse outcomes. Amongst these genes, we note that C5aR1 expression is transiently induced following radiotherapy, likely as a stress response mounted to promote tumour cell survival. Interestingly, in AKPT subcutaneous tumour models, C5aR1 is expressed by both the tumour epithelium and stroma. We find that epithelial expression is more prominent at baseline and early timepoints post irradiation while stromal expression dominates at later timepoints. Increased C5aR1 stromal expression at later timepoints may reflect recruitment of C5aR1 expressing immune cells into the stroma following irradiation (as indicated in **Figure 1C**). Interestingly, in rectal cancer patients we find that those classified as CMS4 have the highest levels of C5aR1 expression compared to the other subtypes. Together, our data indicate that patients with tumours displaying immunosuppressive features and high C5aR1 expression could represent patient populations most likely to benefit from C5aR1 targeting therapy.

Previous reports investigating the effects of complement inhibition on radiotherapy response have focused on the role of complement in modulating anti-tumour immune responses, albeit with conflicting results (Elvington et al., 2014; Surace et al., 2015). Furthermore, none of these reports considered the effect of complement-targeting therapies on cancer-cell intrinsic functions (Bai et al., 2019; Block et al., 2019; Cho et al., 2014). Whether targeting complement would have the same effects in tumour and normal tissues had, until this study, remained completely unexplored. Using a combination of RNA-sequencing and *in silico* mining of depmap, CanSAR, and patient datasets, we identify C5aR1 as a druggable target for enhancing stress-specific cell death in tumours. Our data indicate that C5aR1 negatively regulates apoptosis in cancer cells by modulating cell survival pathways such as NF-κB, and that attenuating such pro-survival signalling can render cancer cells more susceptible to cell death following RT. Importantly, in the normal intestine targeting C5aR1 does not result in increased apoptosis and, in fact, C5aR1 deficiency appears to confer a protective phenotype (which might be explained by the lack of attenuated NF-κB signalling). Why or how C5aR1 may differentially regulate signalling in normal and malignant cells remains to be elucidated. However, divergent consequences of autocrine complement signalling between cell lines have been previously reported (Daugan et al., 2021b). To therapeutically target C5aR1, we used the specific antagonist PMX205 which has FDA and EMA “orphan drug” approval for ALS; allowing accelerated progression to clinical trials (Ricklin and Lambris, 2016). This class of cyclic peptides have minimal penetrance into the cell (Niyonzima et al., 2021) and should therefore primarily impact cell-surface C5aR1 (and its effects on GPCR signalling). This is important since recent reports indicate that C5aR1 intracellular pools are present in tumour cells where they contribute to tumourigenesis through β-catenin stabilisation (Ding et al., 2022).

We have compared, for the first time, single dose, and equivalent fractionation doses in colorectal cancer models to specifically assess the effect of targeting complement at the level of C5aR1. In both settings we observe that PMX205 improves tumour radiation response. We propose that increased PMX205-mediated cell death is a new mechanism that can be exploited to improve radiation responses even in tumours displaying immunosuppressive features. The importance of modulating anti-tumour immune host responses following C5aR1 targeting has been highlighted in previous studies and ourselves and others have observed reduced tumour burden in C5aR1^-/-^ mice (data not shown) (Ding et al., 2022; Markiewski et al., 2008; Surace et al., 2015). The defect in tumour uptake displayed by C5aR1^-/-^ mice complicates the investigation of tumour radiation responses in this model and might have contributed to some of the previous conflicting reports on the effects of targeting the complement system on tumour radiation responses. The reduced tumour CD8^+^ T-cell infiltration observed here, however, is in line with a previous study indicating that local production of anaphylatoxins C3a and C5a and signalling through their respective receptors is required for dendritic cell maturation and CD8^+^ T-cell activation (Surace et al., 2015).

Overall, this study indicates that increased complement gene expression is part of the stress response mounted by irradiated tumours to sustain tumour cell survival via C5aR1 signalling. These data therefore indicate that, beyond its previously described functions, C5aR1 can also sustain tumour cell survival in a cell intrinsic manner. Consequently, targeting complement receptor C5aR1 can improve radiotherapy responses even in tumours displaying reduced CD8+ T-cell infiltration. Importantly, C5aR1’s pro-survival functions appear to be malignant cell specific since increased apoptosis is not observed in the normal intestinal epithelium following C5aR1 loss. This work is relevant since identifying targets that specifically modulate cancer cell radiosensitivity and can do so even in the absence of robust tumour CD8^+^ T cell infiltration could lead the way to improving treatment outcomes for the most difficult to treat tumours.

## Materials and Methods

### Data availability

The RNA-seq data generated for this paper have been deposited at the ArrayExpress database at EMBL-EBI (www.ebi.ac.uk/arrayexpress) and are publicly available as of the date of publication. The data from timecourse study after radiation treatment of heterotopic AKPT organoid-derived tumours are available under accession number E-MTAB-12538, and the data from the intestinal organoids treated with the complement C5a receptor 1 (C5aR1) antagonist PMX205 and/or irradiation are available under accession number E-MTAB-12548.

Publicly available data were analysed in this paper. The analysis of wild-type, AKPT, and KPN intestinal organoids used data deposited at the ArrayExpress database under accession number E-MTAB-11769 (Gil Vazquez et al., 2022).

### Cell lines and treatments

HCT 1 1 6 male adult human epithelial colorectal carcinoma cells originally purchased from ATCC^®^ were used. HT-29 female adult colorectal adenocarcinoma cells were also purchased from ATCC^®^. MC38 murine C57BL6 colon adenocarcinoma cells were a kind gift from Edward Graves. Cells were grown in DMEM with 10% FBS, in a standard humidified incubator at 37°C and 5% CO_2_. All cell lines were routinely tested for mycoplasma and found to be negative.

### AKPT organoid culture

Organoids were sourced from Eoghan Mulholland working within the A:CCSMN. These were generated from male mice with tumours from liver metastasis formed from a tamoxifen-induced colorectal cancer mouse model (with APC ^-/-^, KRAS ^G12D^, p53 ^-/-^, TGFβR1^-/-^). Organoids were grown within 35 μl of Matrigel, DMEM/F12 media mixture (2:1) with 500 μl of media overlaid within each well of a 24-well plate. Overlaid medium was supplemented with epidermal growth factor (EGF) and noggin at a concentration of 100 ng/ml and 50 ng/ml respectively. Organoids were maintained through passaging every three days.

For survival fraction studies, organoids were used 24 hours after passaging and plated into a 24-well plate. Vehicle or PMX205 treatment was carried out as previously described and added into the surrounding media 1 hour prior to irradiation. Organoid irradiation was carried out using an x-ray irradiator with lead shielding allowing for cumulative doses across a single plate (within a twenty-four well plate, two columns of four received 0Gy, two received 4Gy, and two received 9Gy). Organoids were imaged over the course of 3 days using a JuLi stage Real-Time Cell History recorder (NanoEnTek Inc., Korea) by using a 4× objective. Using the images, organoids were manually counted at day 0 and 3 with a surviving fraction subsequently calculated.

### PMX205 treatment

For all *in vitro* experiments cells were treated with 10 μg/ml PMX205 (Tocris #5196) dissolved in 20% ethanol/water. Vehicle control in these experiments refers to 20% ethanol/water. Cells were pre-treated for 1 hour before irradiation.

### Animal studies

MC38 cells (5 × 10^5^) were injected subcutaneously into 6-8 week old female C57/BL6 mice (JAXX) at a single dorsal site. Growing tumours (average, 80-100 mm^3^) were treated with irradiation (either 9 Gy as a single dose or equivalent fractionated doses 3 × 4.45 Gy) and either vehicle or PMX205 for the days flanking the irradiation. In order to perform irradiations of subcutaneous tumours, mice were anesthetised in a knockdown chamber with a mixture of 3% isoflurane and 100% O_2_ and placed inside the irradiator cabinet on the subject stage. Anesthesia was maintained using 1.5% isoflurane in O_2_ delivered via a nose cone. Mice were monitored for breathing with a webcam fitted in the cabinet. The prescribed dose was calculated for a point chosen approximately in the middle of the tumour. The beam angle was chosen so as to target the superficial tumour while sparing critical organs within the mouse. The X-Rad SmART (Precision X-Ray Inc., North Branford, CT) was used for the irradiation of subcutaneous tumours. The X-ray tube and detector plate are mounted on a U shape gantry that rotates 360° in X and Y plane around the animal stage. The animal stage is supplied by a nose cone delivering isoflurane anesthesia, and can move ±10cm in the X and Y directions and ±15cm in the Z direction. Pre-treatment computed tomography (CT) images were acquired to facilitate treatment planning, using a beam energy of 40kVp, a beam filter of 2 mm Al, and a voxel size of 0.2 or 0.1 mm. CT images collected were loaded onto RT_Image and a treatment plan was created using a single beam. The open-source RT_Image software package, version 3.13.1, running on IDL version 8.5.1 was used to visualize CT images and perform treatment planning (Graves et al., 2007). Therapeutic irradiations were performed using an x-ray energy of 225kVp, a current of 13mA, a power of 3000 Watts, and a beam filter of 0.3 mm Cu, producing a dose rate of ~300cGy/min at the isocenter. Treatment x-ray beams were shaped using a 10 or 15 mm collimator to selectively irradiate the target while sparing adjacent tissue when performing xenograft irradiation. For total abdominal irradiations the 20 mm collimator with two beams; 0 and 180 degrees was used. After radiation delivery mice were removed and placed in a recovery box prior to being returned to their cage.

Subcutaneous tumours were measured with the use of callipers and volumes calculated by the ellipsoid estimation method as previously described (Taniguchi et al., 2014). Mice were euthanised as per APLAC guidelines in a CO_2_ chamber and by cervical dislocation.

For heterotopic AKPT organoid-derived tumour models, female C57BL/6 mice were purchased from Charles River at 5-6 weeks of age. Mice were housed within a pathogen-free environment with a 12-hour light cycle at the RRI Biomedical Science Animal Unit at the University of Oxford. AKPT organoids were suspended in a PBS, Matrigel mixture (1:1) prior to subcutaneous injection. Resulting tumours were monitored once a day and their volume determined from: Length × Width × Height × 0.52 using calliper measurements. For experiments shown in Figures 1 and 2, once the mean tumour volume across the mice reached 150-200 mm^3^, mice received irradiation using the Gulmay Medical RS320 irradiator (300kV, 10mA, 1.81Gy/min).

For the experiments shown in Figure 7, four groups of mice (n=10) were established: control, PMX205 treatment, Irradiation, and a combination of irradiation plus PMX205 treatment. Once mean tumour volume for all mice reached 100 mm^3^, treatment was then initiated. PMX205 was delivered by oral gavage once a day for three days. For groups receiving radiation, 9 Gy was delivered using the Gulmay Medical RS320 irradiator (300kV, 10mA, 1.81Gy/min). Once tumour size reached a length by width 1.2 cm geometric mean mice were euthanised by a Schedule 1 method.

For normal tissue studies, total abdominal irradiation of either C57/BL6 (JAXX)(treated with vehicle or PMX205) or WT and C5aR1^-/-^ BALBc/J (JAXX) mice was performed on anaesthetised animals (with the use of ketamine 100mg/kg/xylazine 20mg/kg) and using a 225 kVp cabinet X-ray system filtered with 0.5 mm Cu (at Comparative Medicine Unit, Stanford, CA). Mice were euthanised as per APLAC guidelines in a CO_2_ chamber and by cervical dislocation.

For *in vivo* experiments involving PMX205 treatment, 10 mg/kg PMX205 (Tocris #5196, or synthesized and purified as previously described (Kumar et al., 2020)) was administered to mice orally flanking the irradiation doses. Vehicle control in these experiments refers to 20% ethanol/water.

Experimental procedures were carried out either under a project licence issued by the UK Home Office under the UK Animals (Scientific Procedures) Act of 1986; under approved protocols by the Institutional Animal Care and Use Committee at Medical University of South Carolina; or in accordance with the National Institutes of Health (NIH) guidelines for the use and care of live animals (approved by the Stanford University Institutional Animal Care and Use Committee). Animal Research Reporting of In Vivo Experiments (ARRIVE) guidelines were used.

### Establishment and culture of mouse intestinal organoids

At least 30 mg of mouse small intestine was used for optimal cell yield and organoid establishment. Once obtained, the tissue was transferred into a 50 ml falcon tube with 30 ml of PBS for short term storage on ice until use. In a cell culture hood, tissues were transferred to a 10 cm Petri dish. Using small scissors, the intestinal section was cut open lengthwise as such that the lumen of the intestine was facing up and was then minced into smaller pieces of ~2 mm. Pieces were rinsed twice with cold PBS and transferred into a 50 ml falcon tube filled in with 30 ml of 10% FBS/PBS solution. The tube was shaken forcefully serval times and the intestinal pieces left to settle by gravity to the bottom of the tube. The 10% FBS/PBS solution was replaced with 25 ml of HBSS/EDTA solution in a 500 ml HBSS solution without ca^2+^ and mg^2+^ (Thermo Fisher Scientific). The tube was incubated in 37°C water bath for 10 minutes and shaken forcefully several times every 2-3 minutes. The crypts were released into the supernatant by shaking the tube. The supernatants containing the isolated crypts were collected and the procedure was repeated twice. Each fraction was filtered through a 150 μm cell strainer to remove debris and for crypts enrichment and 1 ml of each fraction was examined under the microscope to check for culturable crypts. 25 ml of 10% FBS/PBS solution was added to the fraction with the highest number of isolated crypts and centrifuged at 1200 rpm at 4°C for five minutes. The pellet was washed with 25 ml of 10% FBS/PBS solution and spined down again. The supernatant was removed carefully and the crypt containing pellets was resuspended in cold Cultrex™ UltiMatrix Reduced Growth Factor (RGF) BME (Bio-Techne). Approximately 250 crypts per 20 ul of BME were seeded in each well of a pre-warmed 48-well plate. The BME was then solidified by a 20-minute incubation in a 37°C and 5% CO_2_ cell culture incubator and overlaid with 500 μl of complete mouse organoid media (1X DMEM/F-12, 1X GlutaMAX, 10 mM HEPES, 10 μg/ml Primocin, 1X N-2 Supplement, 1X B-27 Plus Supplement, 0.1% BSA, 25% R-spondin-1 conditioned medium, 10 ng/ml Mouse EGF, 100 ng/ml Mouse Noggin, 3 μM CHIR 99021) with 10 μM Y-27632 (to ameliorate organoid establishment and survival). Complete media was subsequently refreshed every two days. Organoids were usually passaged every 5–10 days by a mechanical approach using Gentle Cell Dissociation Reagent (GCDR) (STEMCELL™).

### Immunohistochemistry

Formalin-fixed and paraffin-embedded 4 μm sections of C5aR1^-/-^ BALBc/J intestines were dewaxed in Histoclear (10 minutes × 2) followed by rehydration in 100% ethanol (5 minutes × 2), 70% ethanol (5 minutes), and 50% ethanol (5 minutes). Sections were then stained with primary C5aR1 antibody (1:1000, Abcam, Cat: ab59390) using EnVision G2 Doublestain System (Dako) according to the manufacturer’s instructions. The whole section was scanned and analysed using the Aperio CS scanner and ImageScope analysis software (Aperio Technologies, Oxford, UK).

### TUNEL assay

ApopTag^®^ (Millipore #S7100) was used to stain 4 μm formalin-fixed paraffin-embedded small intestinal or AKPT tumour sections. Intestine stained slides were scanned with a NanoZoomer 2.0-RS Digital Slide Scanner (Hamamatsu). For small intestines, the number of TUNEL+ cells per crypt (or crypts and villi) were manually counted in at least 10 fields of view per section. Stained slides for AKPT tumours were scanned with Aperio CS scanner (Aperio Technologies, Oxford, UK) and images were then analysed with QuPath software. The tumour areas within the sections were identified on the basis of the histological structure, and the TUNEL+ areas were then normalised to haematoxylin-positive areas to calculate the percentage of TUNEL+ cancer cells in each section.

Apoptosis assessment by morphology in cell lines was carried out as previously described (Olcina et al., 2015; Olcina et al., 2020). Adherent and detached cells (in media) were collected and fixed in 4% PFA for 15 minutes at room temperature. PFA was then removed, and cells were washed in 1 ml PBS. 10 μl of fixed cells in PBS was placed in each slide and mixed with ProLong^™^ Gold antifade mountant with DAPI (Invitrogen #P36935). A coverslip was placed on top of the sample and slides were left to dry overnight. Slides were imaged using a DSM6000, DMi8 or DMI6000 (Leica) microscope or Nikon Ni-E with 40× or 60× oil objectives. The number of cells with fragmented DNA and the total number of cells per field was counted (with typically at least 10 fields counted for every treatment).

### Multiplex and HALO analysis

Multiplex immunofluorescence staining of AKPT tumours was performed on 4 μm thick FFPE sections using the OPAL protocol (Akoya Biosciences, Marlborough, MA). The Leica BOND RXm autostainer (Leica Microsystems, Wetzlar, Germany) was used to conduct this staining. Staining cycles were conducted six consecutive times using the following primary antibody-Opal fluorophore pairs:

#### Immune panel

(1) Ly6G (1:300, 551459; BD Pharmingen)–Opal 540; (2) CD4 (1:500, ab183685; Abcam)–Opal 520; (3) CD8 (1:800, 98941; Cell Signaling)– Opal 570; (4) CD68 (1:1200, ab125212; Abcam)–Opal 620; (5) FoxP3 (1:400, 126553; Cell Signaling)–Opal 650; and (6) E-cadherin (1:500, 3195; Cell Signaling)–Opal 690.

#### Stroma panel

(1) Gremlin 1 (1:750, AF956; R&D)–Opal 540; (2) CD34 (1:3000, ab81289; Abcam)–Opal 520; (3) CD146 (1:500, ab75769; Abcam)–Opal 570; (4) SMA (1:1000, ab5694; Abcam)–Opal 620; (5) Periostin (1:1000, ab227049; Abcam)–Opal 690; and (6) E-cadherin (1:500, 3195; Cell Signaling)–Opal 650.

Tissues sections were incubated for 1 h in primary antibodies and detected using the BOND Polymer Refine Detection System (DS9800; Leica Biosystems, Buffalo Grove, IL) in accordance with the manufacturer’s instructions, substituting DAB for the Opal fluorophores, with a 10-min incubation time and withholding the hematoxylin step. Antigen retrieval at 100°C for 20 min, in accordance with standard Leica protocol, with Epitope Retrieval Solution one or two was performed prior to each primary antibody being applied. Sections were then incubated for 10 min with spectral DAPI (FP1490, Akoya Biosciences) and the slides mounted with VECTASHIELD Vibrance Antifade Mounting Medium (H-1700-10; Vector Laboratories). Whole slide scans and multispectral images (MSI) were obtained on the Akoya Biosciences Vectra Polaris. Batch analysis of the MSIs from each case was performed with the inForm 2.4.8 software provided. Finally, batched analysed MSIs were fused in HALO (Indica Labs) to produce a spectrally unmixed reconstructed whole-tissue image. Cell density analysis was subsequently performed for each cell phenotype across the three MPIF panels using HALO. HALO Image Analysis Platform version 3.5.3577 and HALO AI version 3.5.3577 (Indica Labs, Inc.) were used. Analysis modules “Area Quntification 2.4.2”, “HighPlex FL 4.2.3” and “Random Forest Classifier” were used.

Cover slips were lifted post multiplex staining and C5aR1 (1:1000, Abcam, Cat: ab59390) antibody was stained for chromogenically on the Leica BOND autostainer. Antigen retrieval was carried out at 100°C for 20 min with Epitope Retrieval Solution 2. Primary antibody incubation at 1:250 dilution for 30 minutes was followed by detection using the BOND™ Polymer Refine Detection System (DS9800, Leica Biosystems) as per manufacturer’s instructions.

### Immunoblotting

Cells were lysed in UTB (9 M urea, 75 mM Tris-HCl pH 7.5 and 0.15 M β-mercaptoethanol) and sonicated briefly before quantification as described in detail in (https://data.mendeley.com).

For intestinal organoid experiments, after the treatment, organoids were harvested by incubation in Corning™ Cell Recovery Solution at 4°C for 30 minutes. Samples were then washed with cold PBS, pelleted (2,000 rpm, 3 min, 4 °C), and lysed immediately in RIPA buffer (Sigma-Aldrich) containing 1:100 protease inhibitor cocktail (Sigma-Aldrich) and 1:100 phosphatase inhibitor cocktail 2 (Sigma-Aldrich). Proteins were quantified with Pierce BCA Protein Assay Kit (Thermo Scientific), using a BSA standard curve (20 to 2000 μg/mL).

4–20% polyacrylamide gels (Bio-Rad) were used for protein separation. Proteins were blotted onto a nitrocellulose membrane (Bio-Rad). The BioRad Chemidoc XRS system or LI-COR Odyssey imaging system was used. In each case experiments were carried out in triplicate and a representative blot is shown unless otherwise stated.

Antibodies used were: β-actin (Sigma # A5441, concentration: 1:5000). hFAB™ Rhodamine Anti-Tubulin (Bio-Rad #12004166), RelA/p65 (Cell Signaling #3034, concentration 1:1000), pRelA/p65-S536 (Cell Signaling #3033T, concentration 1:1000), total AKT (Cell Signalling #2920, concentration 1:1000), AKT-T308 (Cell Signalling #13038, concentration 1:1000), p44/p42 (Cell Signalling #4696, concentration 1:1000), phospho p44/p42 (Cell Signalling #4376), γH2AX (Millipore #05-636-1, concentration 1:500).

### Reverse Phase Protein Array (RPPA)

RPPA was performed by the University of Texas MD Anderson RPPA core as described in the published protocol (https://www.mdanderson.org/research/research-resources/core-facilities/functional-proteomics-rppa-core.html). For statistical analysis on differently expressed proteins, linearised (standard curve) normalised (to protein loading) relative protein levels were analysed using two-tailed Student’s *t* test (t.test()) in R.

### Flow cytometry analyses

For analysis of immune populations in tumour draining lymph nodes, 5 × 10^5^ MC38 cells were injected into the flank of 8-week-old female C57/BL6 mice. When tumours were approximately 50 mm^3^ in volume, the mice were treated via oral gavage with either vehicle or PMX205 (10 mg/kg) 24 hours before irradiation. These treatments were repeated 1-hour prior irradiation.

For the radiation therapy, the tumour-bearing mice were anesthetised with ketamine/xylazine (80mg/Kg/ 10mg/Kg) and a dose of 9 Gy was delivered to the tumour site using a Varian TrueBeam 2041 clinical linear accelerator. 24 hours after irradiation the treatment with vehicle or PMX205 was repeated. One week after irradiation, the mice were sacrificed and tumour draining lymph nodes were collected for FACS analysis. For flow cytometry staining, cells were incubated with anti-FcR antibody (clone 24G2) and stained with the following surface antibodies: anti-CD45.2 (clone 104, eBioscience), CD3 (clone 17A2, eBioscience), CD4 (cloneRM4-5), CD8a (53-6-7), B220 (clone RA3-6B2), CD25 (clone PC61), a-CD49b (clone DX5), anti-IFN-γ (clone XMG1.2), anti-FoxP3 (clone FJK-16s eBioscience), anti-TNF-α (clone MP6-XT22), anti-Granzyme B (clone BG11, BioLegend). Dead cells were stained using a Fixable Viability Dye eFluor 506 (eBioscience). For cytokine staining, cells were stimulated for 4 hours with 50ng/ml PMA and 500ng/ml Ionomycin (Sigma-Aldrich), washed, and after surface staining, permeabilised with Cytofix/Cytoperm buffer according to manufacturer instructions. For Foxp3 staining the permeabilisation was performed using the FoxP3 permeabilisation buffer (eBioscience). All antibodies were purchased from BD Pharmingen unless otherwise specified. Samples were acquired with Fortessa LSR (BD Bioscience) and analyzed with FlowJo software (TreeStar).

For analysis of tumour immune infiltration, female C57/BL6 MC38 tumour bearing mice were euthanised as per APLAC guidelines. Tumour tissue was digested into a single suspension using the murine tumour dissociation kit from Miltenyi Biotech (Auburn, CA) as per the manufacturer’s protocol. After RBC lysis, cells were re-suspended in PBS, counted and then stained with Zombie NIR (BioLegend, San Diego, CA) for live/dead cell discrimination. Nonspecific binding was blocked using an anti-mouse CD16/32 (BioLegend, San Diego, CA) antibody. Following which, cell surface staining was performed using fluorophore conjugated anti-mouse CD45.1 (30-F11), CD11b (M1/70), CD11c (N418), Ly6G (1A8), Ly6C (HK1.4), CD8 (53-6.7), from Biolegend (San Diego, CA). Flow cytometry was performed on LSR Fortessa (BD Biosciences) in the Radiation Oncology Dept. FACS facility. Cell acquisition was performed the following day with FACSDiva software on an LSR II flow cytometer (BD Biosciences, San Jose, CA) and analysed with FlowJo software (Tree Star Inc., San Carlos, CA). Compensations were attained using Anti-Rat and anti-hamster compensation beads (BD Biosciences). For fixable live/dead staining, compensation was performed using ArC amine reactive compensation beads (BD Biosciences).

Gating schemes of dissociated tissues was as follows: Immune cells- (ZNIR-CD45+), CD8 T-cells (ZNIR-CD45+CD8+), Neutrophils (ZNIR-CD45+CD11b+Ly6G+), Monocytes (ZNIR-CD45+CD11b+Ly6C+).

### Cell cycle analysis

Cells were harvested with trypsin, washed in PBS and fixed in 1 ml of 70% ethanol. Fixed cells were incubated on ice for 30 minutes. Following incubation, cells were pelleted and washed twice in PBS. Cell pellets were treated with 50 μL of PureLink™ RNase A (invitrogen, 12091-021, 100μg/ml) before adding 400 μL of propidium iodide solution (invitrogen, 00-6990-50, 50μg/ml) in PBS. Cells were incubated at room temperature for 10 minutes. Following incubation, the cell cycle was analysed with CytoFLEX Flow Cytometer (Beckman Coulter, Inc.) and data analysis was performed using FlowJo Software (version 10.7.2).

### qRT-PCR

RNA was extracted using Trizol (Invitrogen/Life Technologies,# 15596018). iScript cDNA synthesis kit (Bio-Rad, #1708891) or Verso cDNA Synthesis kit (thermoscientific, #AB-1453/B) was used to reverse transcribe cDNA from total RNA according to manufacturer’s instructions. Relative mRNA levels were calculated using the dCt methodology using a 7900HT Fast Real-Time PCR System. Primers used: ACTB F: ACATCCGCAAAGACCTCTACG, ACTB R: TTGCTGATCCACATCTGCTGG; C5AR1 F: TCCTTCAATTATACCACCCCTGA, C5AR1 R: ACGCAGCGTGTTAGAAGTTTTAT; BCL2 F: TTGCCAGCCGGAACCTATG BCL2 R: CGAAGGCGACCAGCAATGATA; C5 F: CTCCTCAGGCCATGTTCATT; C5 R: TCTTTTGGCTGGCTTCAAGT; 18s F: GTGGAGCGATTTGTCTGGTT, 18s R: ACGCTGAGCCAGTCAGTGTA; C5ar1 F: ACATGGACCCCATAGATAAC: C5ar1 R: ACCACCGAGTAGATGATAAG; c5 F: TACCAATGCCAACCTGGTGAAAGG, c5 R: TCTGCAGAACCTCTTTGCCCATGA; Bcl2 F: GTCGCTACCGTCGTGACTTC, Bcl2 R: CAGACATGCACCTACCCAGC, Bcl2l1 F: GACAAGGAGATGCAGGTATTGG, bcl2l1 R: TCCCGTAGAGATCCACAAAAGT, Xiap1 F: CGAGCTGGGTTTCTTTATACCG, Xiap1 R: GCAATTTGGGGATATTCTCCTGT

### siRNA transfection

NFKBIA (L-004765-00), RelA (L-003533-00), C5aR1 (L-005442-00) or non-targeting RNAi negative control (Scramble, D-001810-10) (all from Dharmacon) were transfected into HCT 116 cells using Lipofectamine^®^ RNAiMax transfection reagent (Invitrogen, #13778075) at a final concentration of 50 nM; according to the manufacturers’ instructions. Cells were harvested 72 hours post-transfection.

### *In silico* screen

Data was accessed through the Depmap portal (https://depmap.org from 10^th^ September 2020). Essentiality scores were calculated by dividing the number of dependent cell lines for each gene/total number of cell lines in either CRISPR-cas9 and RNAi screens. To identify stress- and cancer-specific dependencies, we applied stringent criteria and looked for genes that were deemed non-essential across all cell lines (genes that when knocked-out/down under baseline conditions do not alter cell survival) while still being expressed across cell lines. Genes described as complement system components, receptors, proteases and regulators (as reported in (Ricklin et al., 2010)) were queried. The calculated essentiality score was presented as a colour in the heatmap. Green = No cell line is dependent on that gene and therefore the gene is not essential. Red = The majority of cell lines are dependent, or the gene is essential. Yellow/Orange = intermediate dependence or essentiality. The essentiality scale is shown at the bottom, left hand side of the heatmap. *ATR* was included in the screen a positive control. Target tractability was assessed by accessing canSAR data as displayed on the Depmap portal. A gene was only considered a hit if it was “druggable” based on structural and ligand-based assessment. Genes that are hits and are druggable are shown in green. Red = not druggable based on structural or ligand-based assessment. C1R and CF1 were not included in the heatmap due to lack of CRISPR-Cas9 or RNAi data. C4A, C4B, VSIG4, C8A, C8B, CD93 and CR1 were not included due to their very low expression in the majority of tissues.

### RNA-sequencing

Transcriptomic profiling of mouse tumour tissues was carried out by 3’RNAseq. Extracted RNA was quantified using RiboGreen (Invitrogen) on the FLUOstar OPTIMA plate reader (BMG Labtech) and the size profiles and integrity were analysed on the TapeStation (Agilent, RNA ScreenTape). Libraries were prepared using the Lexogen QuantSeq 3’ mRNA-Seq kit FWD kit (Lexogen, Cat no. 015.2 × 96) and the Lexogen UMI Second strand synthesis module for QuantSeq FWD (Lexogen, Cat No. 81.96) following the manufacturer’s instructions. Individual libraries were quantified using a Qubit Fluorometer (Invitrogen), and the size profiles were analysed on the Agilent TapeStation. Individual libraries were normalised and pooled accordingly to multiplex for sequencing. Libraries were sequenced on a NextSeq 500 instrument (Illumina) as 75-bp single-end reads.

Raw sequence reads were subjected to adapter trimming using *BBduk* (BBTools *ver. 38.46*) Trimmed reads were aligned to the Genome Reference Consortium mouse genome build 38 (GRCm38) build of the mouse reference using *STAR* (ver. *2.7.0f*). Ensembl 96 annotations were used for alignment and subsequent quantifications. Gene expression was quantified using *featureCounts* (ver. 1.6.4). Further analysis of RNA-seq data were carried out in the R statistical environment (*ver. 4.0.3*). Differential expression analyses were performed using the *limma* (*ver. 3.46*) package. Gene Set Enrichment Analyses were performed using the *fgsea* package (*ver. 1.16*). Network graphs were drawn from the significantly-enriched pathways (*P_FDR_* ≤ 0.10) with the *PPInfer* package (*ver. 1.16.0*) using the Fruchterman-Reingold algorithm for visualisation.

### Organoids RNA extraction for sequencing

After the treatment, organoids were harvested by direct incubation in 350 μl RLT plus lysis buffer (Qiagen RNeasy^®^ Plus Micro kit) in each well for 5 minutes and their RNA was extracted using the Qiagen RNeasy^®^ Plus Micro kit according to the manufacturer protocol. RNA quality and concentration were measured with Agilent Bioanalyzer 2100 (Agilent) and Nanodrop (Thermo Fisher Scientific) respectively. RNA library preparation, sequencing, and data analysis were outsourced to Novogene (UK).

### CMS in rectal tumours

The Grampian cohort profiled by the S:CORT consortium was used. Pre-treatment rectal tumour biopsies from patients treated with radiotherapy and capecitabine were selected (N=129). Xcell array data was normalised and CMS called using the R tool CMScaller (Eide et al., 2017). Patients gave consent to biobank their samples in the Grampian Biorepository (ref no TR000028, 1/10/2014) with release of linked anonymised clinical data for ethically approved research projects.

This study involves human subjects, and all samples were obtained following individual informed consent for research projects and subsequent ethical approval by the National Research Ethics Service in the UK (REC ref 15/EE/0241, IRAS ref 169363).

## Author contributions

Methodology C.B., D.ML., D.M., S.M., D.K.M., R.K.K. S.G., G.N.V., M.D.D., T.S., A.E., D.J., J.Y., E.D., and M.M.O. Writing original draft, M.M.O. Writing review and editing, M.M.O., D.ML., A.J.G., D.K.N., S.G. G.N.V., M.D.D., A.E., E.D., E.M., J.Y., S.J.A.B., M.S., S.J.L., Q.T.L., and T.M.W. Conceptualisation, M.M.O and A.J.G. Investigation, C.B., D.ML., D.M., S.M., D.K.N., R.K.K., S.G., K.C. G.N.V., T.S., M.D.D., E.D., Y.J., E.J.M., D.J., and M.M.O. Formal analysis, M.M.O., D.K.N., R.K.K., C.B., D.ML., D.M., E.D., D.J., S.G., and G.N.V. Resources, M.M.O, A.J.G., T.M.W., S.J.A.B., S.J.L, Q.T.L, E.G.G., S.G., A.C.K., and M.S. Supervision M.M.O., A.J.G., S.J.L., T.M., Q.T.L. Funding acquisition, M.M.O., A.J.G., S.J.L., and M.S.

## Acknowledgements

This work was supported by MRC (MC_UU_00001/10), NIH Grants CA-67166 and CA-197713, the Silicon Valley Foundation, the Sydney Frank Foundation and the Kimmelman Fund (AJG). This work was also supported by Cancer Research UK (CR-UK) grant number C5255/A18085, through the Cancer Research UK Oxford Centre. MMO was a Cancer Research Institute Irvington Fellow supported by the Cancer Research Institute and Stiftung für Forschung, Medical Faculty, UZH. C.B is supported by an International Accelerator Award, ACRCelerate funded by Cancer Research UK (A26825 and A28223). RKK was supported by Stanford ChEM-H Undergraduate Scholars Program. MS and MMO were supported by a project grant from the Swiss National Foundation (31003A_163141). ED is supported by the S:CORT Consortium which is funded by a grant from the Medical Research Council and Cancer Research UK. TMW is supported by a fellowship from the National Health and Medical Research Council (2009957). We would also like to acknowledge the Tissue Histopathology Laboratory (University of Oxford) and, in particular, Leticia Campo Urriza and Molly Browne.

