## Supplemental Figures for "Innate immune receptor C5aR1 regulates cancer cell fate and can be targeted to improve radiotherapy in tumours with immunosuppressive microenvironments"

+ Joint author.

*Corresponding Author:

Monica M. Olcina
Current address:
Oxford Institute of Radiation Oncology
University of Oxford, Old Road Campus Research Building
Roosevelt Drive, Oxford, OX37DQ.

**Running tittle:** Targeting C5aR1 to improve radiotherapy in immune excluded tumours

**Keywords:** C5aR1, colorectal cancer, radiotherapy, complement

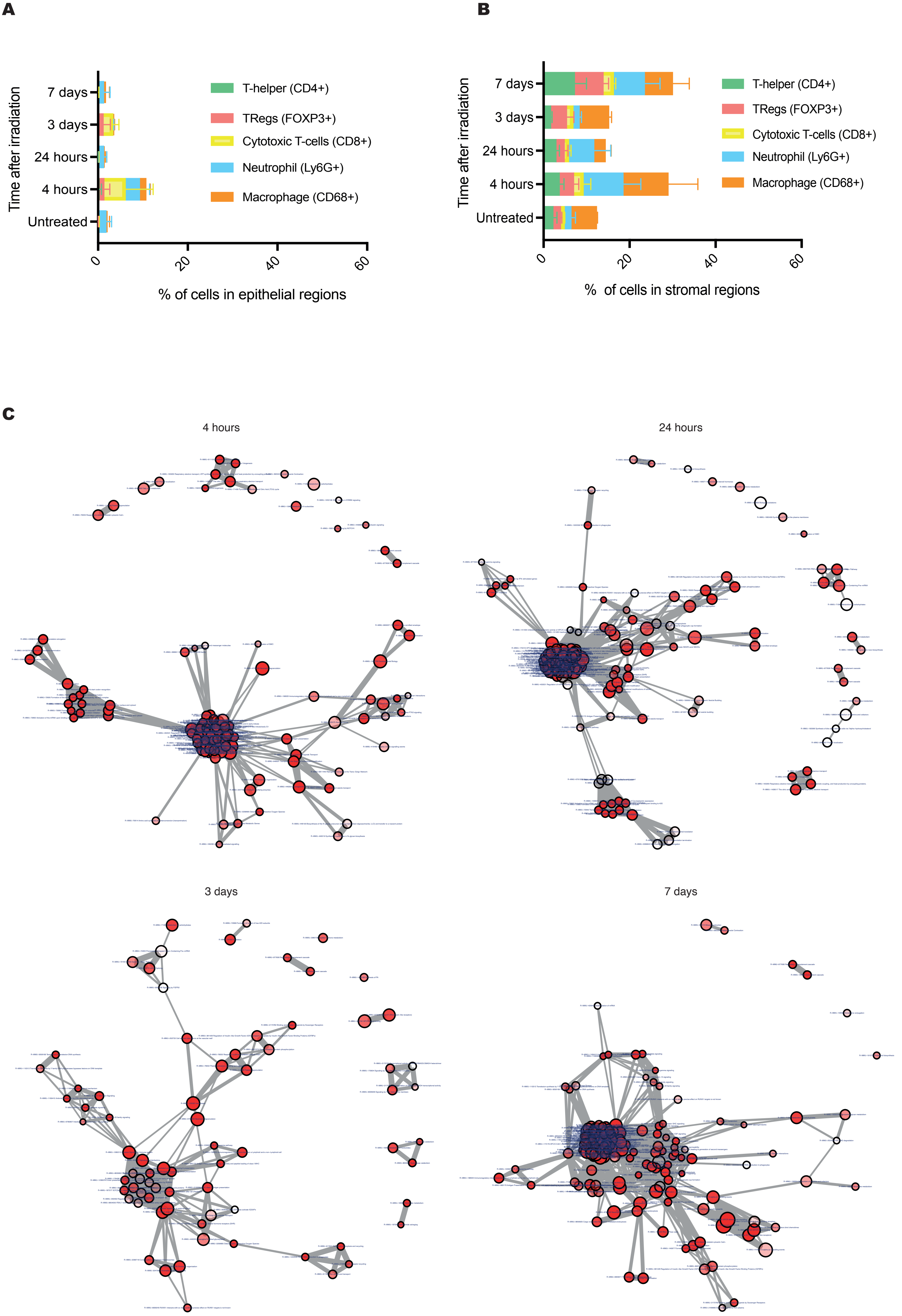
A
Time after irradiation
7 days
3 days
24 hours
4 hours
Untreated
0
20
40
60
% of cells in epithelial regions
T-helper (CD4+)
TRegs (FOXP3+)
Cytotoxic T-cells (CD8+)
Neutrophil (Ly6G+)
Macrophage (CD68+)
B
Time after irradiation
7 days
3 days
24 hours
4 hours
Untreated
0
20
40
60
% of cells in stromal regions
T-helper (CD4+)
TRegs (FOXP3+)
Cytotoxic T-cells (CD8+)
Neutrophil (Ly6G+)
Macrophage (CD68+)
C
4 hours
24 hours
3 days
7 days

**Supplementary Figure 1. Identification of radiation-responsive targets in immunosuppressive tumours**

- **(A)** Machine learning-based quantification of immune cell infiltration in epithelial regions following multiplex staining at different timepoints following irradiation.
- **(B)** Machine learning-based quantification of immune cell infiltration in stromal regions following multiplex staining at different timepoints following irradiation.
- **(C)** Network graphs of mouse Reactome pathways found significantly enriched ($P_{FDR} \leq 0.10$) after Gene Set Enrichment Analysis**.** Significantly-enriched Reactome pathways for each timepoint are shown as network graphs using the Fruchterman-Reingold algorithm, as in Figure 1D and E. The network graphs are shown here in a larger format, with the nodes labelled with their Reactome pathway names. Each node is a pathway gene set, with the size of the nodes being proportional to the size of the gene set and pathways with lower $P_{FDR}$ appearing as less transparent nodes. Connections between nodes depend on the proportion of overlapping genes between two gene sets.

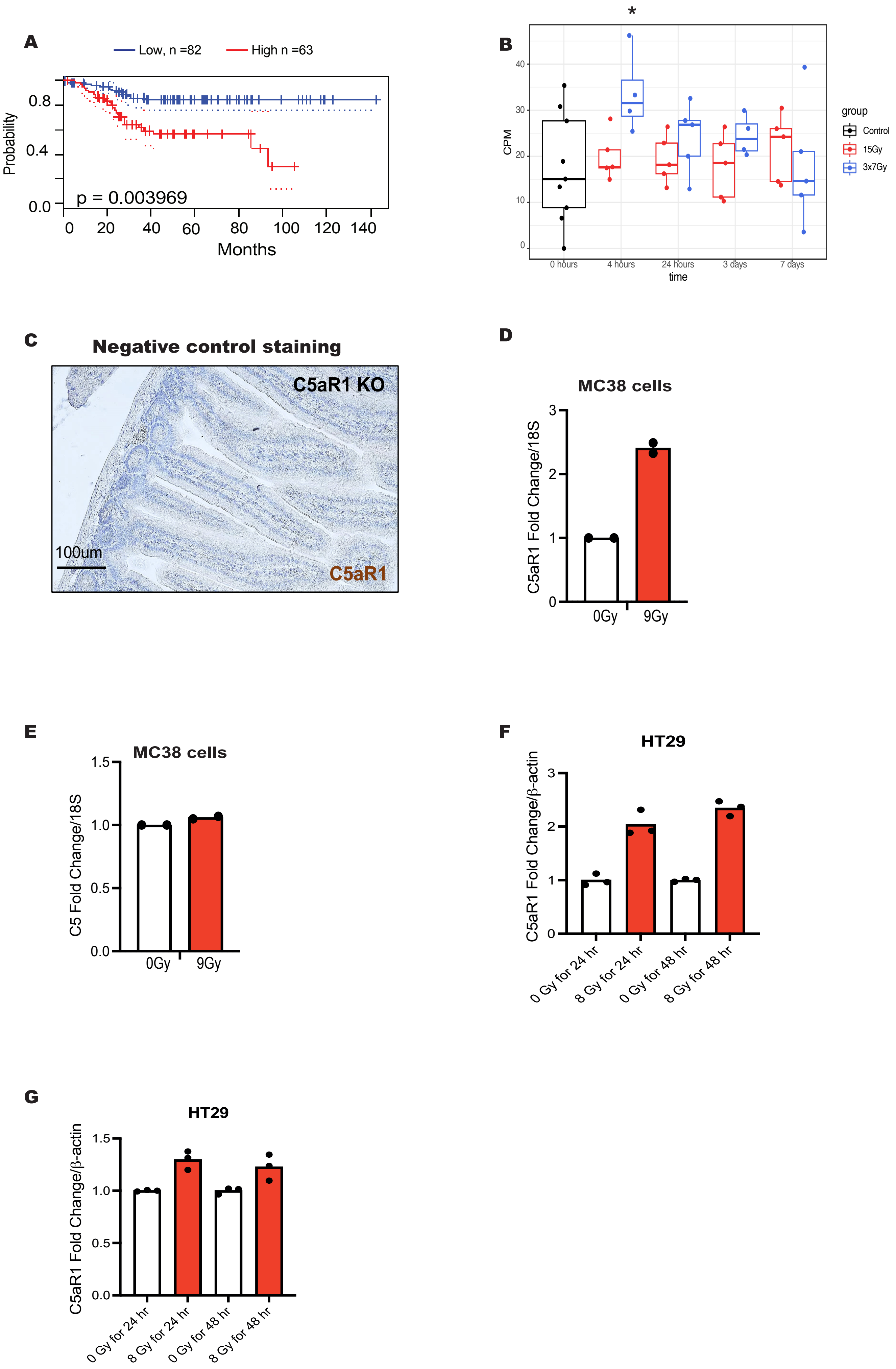
A
Low, n =82
High n =63
Probability
0.8
0.4
0.0
p = 0.003969
0
20
40
60
80
100
120
140
Months
B
*
CPM
40
30
20
10
0
0 hours
4 hours
24 hours
3 days
7 days
time
group
Control
15Gy
3x7Gy
C
Negative control staining
C5aR1 KO
100um
C5aR1
D
MC38 cells
C5aR1 Fold Change/18S
3
2
1
0
0Gy
9Gy
E
MC38 cells
C5 Fold Change/18S
1.5
1.0
0.5
0.0
0Gy
9Gy
F
HT29
C5aR1 Fold Change/β-actin
3
2
1
0
0 Gy for 24 hr
8 Gy for 24 hr
0 Gy for 48 hr
8 Gy for 48 hr
G
HT29
C5aR1 Fold Change/β-actin
1.5
1.0
0.5
0.0
0 Gy for 24 hr
8 Gy for 24 hr
0 Gy for 48 hr
8 Gy for 48 hr

**Supplementary Figure 2: C5aR1 is a radiation-responsive druggable target**

- **(A)** Prognoscan KM curve for colorectal cancer patients with high (red) or low (blue) C5AR1 mRNA expression levels is shown. This analysis was based on the PrognoScan database (http://www.prognoscan.org/) using the publicly available Gene Expression Omnibus (http://www.ncbi.nlm.nih.gov/geo).
- **(B)** C5aR1 expression following RNA-seq of AKPT tumour receiving either 15 Gy or 3 x 7 Gy irradiation.
- **(C)** Immunohistochemistry staining of small intestine section of a C5aR1$^{-/-}$ mouse used as a negative control for the C5aR1 staining shown in Figure 2F.
- **(D)** mRNA expression of *C5AR1/housekeeping* in MC38 cells treated with either 0 or 9 Gy irradiation. Individual points indicate average from biologically independent replicates. n=2.
- **(E)** mRNA expression of *C5AR1/housekeeping* in MC38 cells treated with either 0 or 9 Gy irradiation. Individual points indicate average from biologically independent replicates. n=2.
- **(F)** mRNA expression of *C5AR1/housekeeping* in HT29 cells treated with either 0 or 9 Gy irradiation. n=1.Individual points indicate technical replicates.
- **(G)** mRNA expression of *C5/housekeeping* in HT29 cells treated with either 0 or 9 Gy irradiation. n=1. Individual points indicate technical replicates.

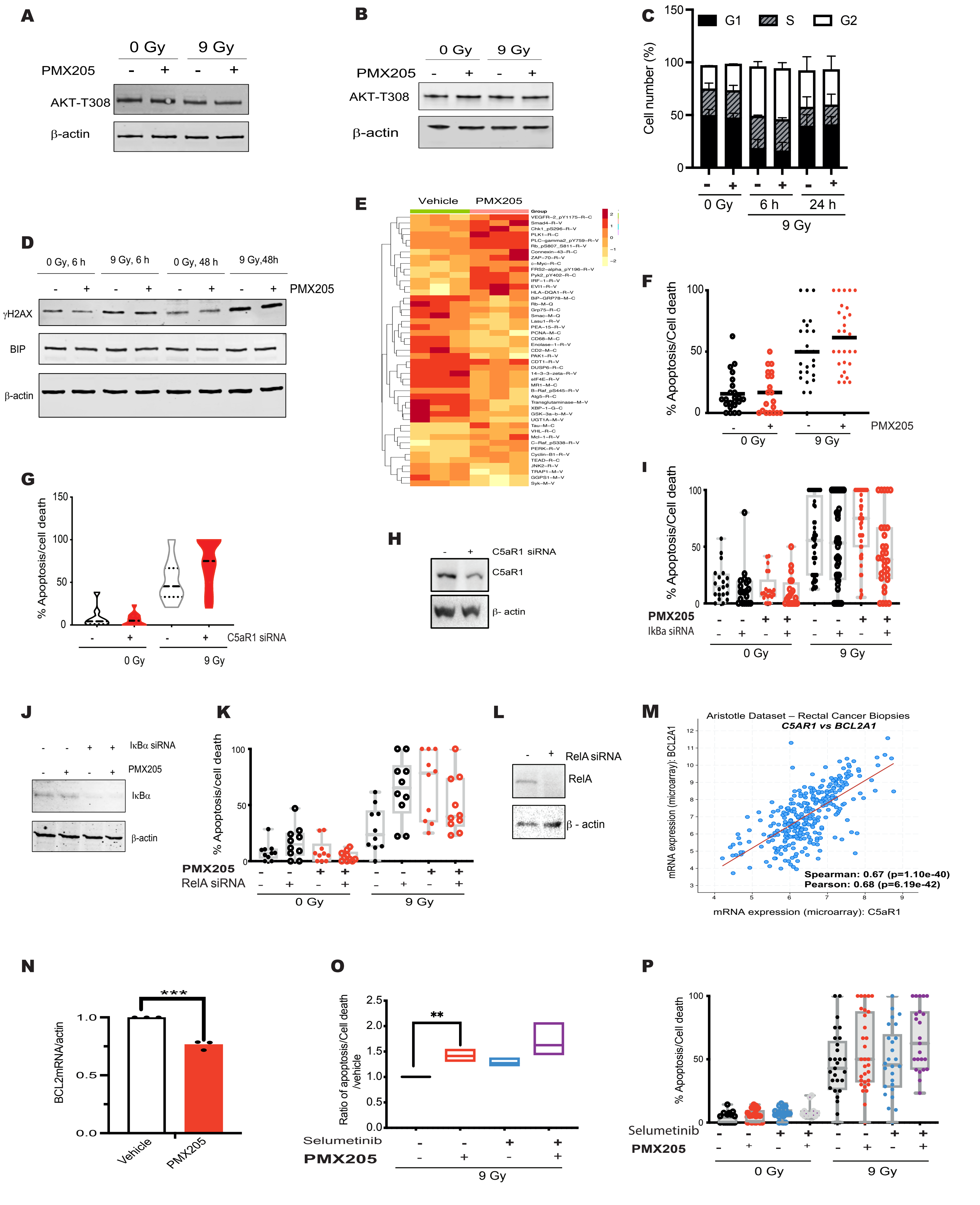
A
0 Gy
9 Gy
PMX205
AKT-T308
β-actin
B
0 Gy
9 Gy
PMX205
AKT-T308
β-actin
C
G1
S
G2
Cell number (%)
0 Gy
6 h
24 h
9 Gy
D
0 Gy, 6 h
9 Gy, 6 h
0 Gy, 48 h
9 Gy,48h
PMX205
γH2AX
BIP
β-actin
E
Vehicle
PMX205
F
% Apoptosis/Cell death
PMX205
0 Gy
9 Gy
G
% Apoptosis/cell death
C5aR1 siRNA
0 Gy
9 Gy
H
C5aR1 siRNA
C5aR1
β- actin
I
% Apoptosis/Cell death
PMX205
IkBa siRNA
0 Gy
9 Gy
J
IκBα siRNA
PMX205
IκBα
β-actin
K
% Apoptosis/cell death
PMX205
RelA siRNA
0 Gy
9 Gy
L
RelA siRNA
RelA
β - actin
M
Aristotle Dataset – Rectal Cancer Biopsies
C5AR1 vs BCL2A1
mRNA expression (microarray): BCL2A1
Spearman: 0.67 (p=1.10e-40)
Pearson: 0.68 (p=6.19e-42)
mRNA expression (microarray): C5aR1
N
***
BCL2mRNA/actin
Vehicle
PMX205
O
**
Ratio of apoptosis/Cell death /vehicle
Selumetinib
PMX205
9 Gy
P
% Apoptosis/Cell death
Selumetinib
PMX205
0 Gy
9 Gy

**Supplementary Figure 3: C5aR1 regulates tumour cell survival under stress**

- **(A)** HT29 cells were treated with 0 or 9 Gy and either vehicle or PMX205 1 hour before irradiation. Cells were harvested 48 hours post-IR. Western blotting was carried with the antibodies indicated. β-actin was used as the loading control.
- **(B)** HCT116 cells were treated with 0 or 9 Gy and either vehicle or PMX205 1 hour before irradiation. Cells were harvested 48 hours post-IR. Western blotting was carried with the antibodies indicated. β-actin was used as the loading control.
- **(C)** Percentage of cells in each cell cycle phase following the same treatments described in (B). n=3.
- **(D)** HCT116 cells were treated with 0 or 9 Gy and either vehicle or PMX205 1 hour before irradiation. Cells were harvested either 6 or 48 hours post-IR. Western blotting was carried with the antibodies indicated. β-actin was used as the loading control.
- **(E)** Heatmap of proteins differentially expressed following reverse phase protein array (RPPA) analysis in HCT116 cells treated with vehicle or PMX205. n=3.
- **(F)** The graph represents the number of dead/apoptotic cells expressed as a % of the whole population for HCT116 cells treated with either vehicle or PMX205 for 1 hour before irradiation (IR) with either 0 or 9 Gy. Cells were harvested 48 hours post-IR. n=3.
- **(G)** The graph represents the number of apoptotic/non-apoptotic cells expressed as a % of the whole population for HCT116 cells treated with either Scr or C5aR1 siRNA and either 0 or 9 Gy IR. Cells were harvested 48 hours post-IR. n=2.
- **(H)** HCT116 cells treated with either Scr or C5aR1 siRNA. Western blotting was carried with the antibodies indicated. β-actin was used as the loading control.
- **(I)** The graph represents the number of apoptotic/non-apoptotic cells expressed as a % of the whole population for HCT116 cells treated with either Scr or IκBα siRNA and either 0 or 9 Gy IR. Cells were harvested 48 hours post-IR.
- **(J)** HCT116 cells treated with 9 Gy and either Scr, IκBα siRNA and vehicle or PMX205. Western blotting was carried with the antibodies indicated. β-actin was used as the loading control.
- **(K)** The graph represents the number of dead/apoptotic cells expressed as a % of the whole population for HCT 116 cells transfected with either Scr or RelA siRNA and treated with either vehicle or PMX205 for 1 hour before irradiation with either 0 or 9 Gy IR. Cells were harvested 48 hours post-IR. Independent fields of view from a representative experiment are show, n=2.
- **(L)** HCT116 cells treated with either Scr or RelA siRNA. Western blotting was carried with the antibodies indicated. β-actin was used as the loading control.

**(M)** mRNA expression of *BCL2/housekeeping* in HCT116 cells treated with either vehicle or PMX205 and 0 or 9 Gy irradiation. n=3.

**(N)** The graph represents the ratio of dead/apoptotic cells expressed as a % of the whole population relative to those cells treated with vehicle (9 Gy). Cells were harvested 48 hours post-IR. n=3. Counted fields of view from all experiments are show, n=3.

**(O)** The graph represents the number of dead/apoptotic cells expressed as a % of the whole population for HT29 cells treated with either PMX205, Selumetinib or a combination of PMX205 and selumetinib and either 0 or 9 Gy IR. Cells were harvested 48 hours post-IR. n=3. Counted fields of view from all experiments are show, n=3.

**A**

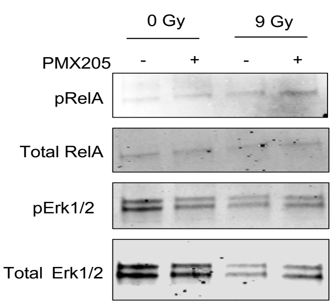

0 Gy
9 Gy
PMX205
-
+
-
+
pRelA
Total RelA
pErk1/2
Total Erk1/2


**B**

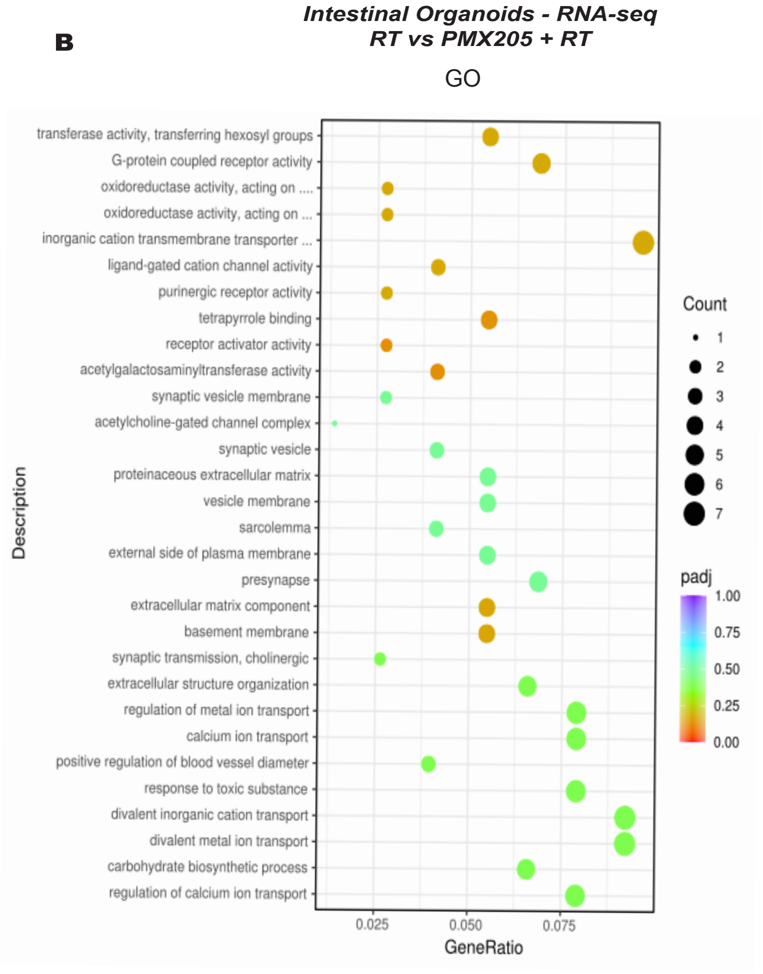

*Intestinal Organoids - RNA-seq*
*RT vs PMX205 + RT*
GO
transferase activity, transferring hexosyl groups
G-protein coupled receptor activity
oxidoreductase activity, acting on ....
oxidoreductase activity, acting on ...
inorganic cation transmembrane transporter ...
ligand-gated cation channel activity
purinergic receptor activity
tetrapyrrole binding
receptor activator activity
acetylgalactosaminyltransferase activity
synaptic vesicle membrane
acetylcholine-gated channel complex
synaptic vesicle
proteinaceous extracellular matrix
vesicle membrane
sarcolemma
external side of plasma membrane
presynapse
extracellular matrix component
basement membrane
synaptic transmission, cholinergic
extracellular structure organization
regulation of metal ion transport
calcium ion transport
positive regulation of blood vessel diameter
response to toxic substance
divalent inorganic cation transport
divalent metal ion transport
carbohydrate biosynthetic process
regulation of calcium ion transport
Description
0.025
0.050
0.075
GeneRatio
Count
1
2
3
4
5
6
7
padj
1.00
0.75
0.50
0.25
0.00


**C**

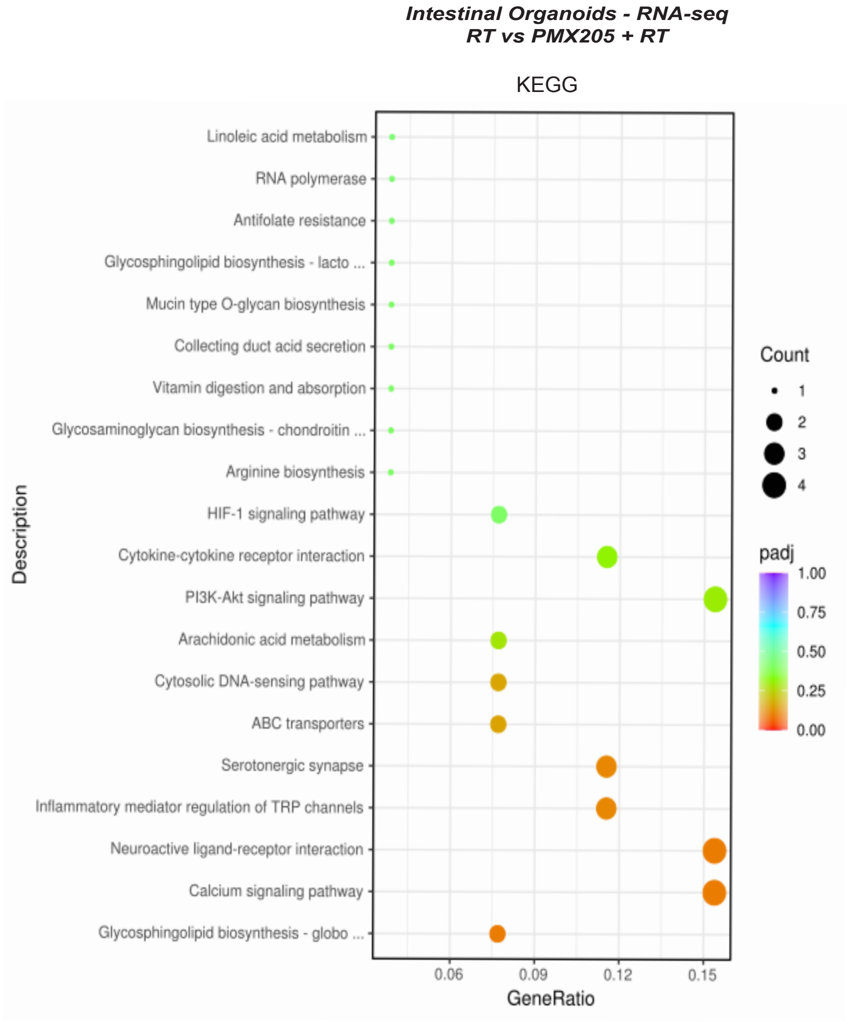

*Intestinal Organoids - RNA-seq*
*RT vs PMX205 + RT*
KEGG
Linoleic acid metabolism
RNA polymerase
Antifolate resistance
Glycosphingolipid biosynthesis - lacto ...
Mucin type O-glycan biosynthesis
Collecting duct acid secretion
Vitamin digestion and absorption
Glycosaminoglycan biosynthesis - chondroitin ...
Arginine biosynthesis
HIF-1 signaling pathway
Cytokine-cytokine receptor interaction
PI3K-Akt signaling pathway
Arachidonic acid metabolism
Cytosolic DNA-sensing pathway
ABC transporters
Serotonergic synapse
Inflammatory mediator regulation of TRP channels
Neuroactive ligand-receptor interaction
Calcium signaling pathway
Glycosphingolipid biosynthesis - globo ...
Description
0.06
0.09
0.12
0.15
GeneRatio
Count
1
2
3
4
padj
1.00
0.75
0.50
0.25
0.00

**Supplementary Figure 4: C5aR1 deficiency does not result in increased apoptosis in healthy intestinal epithelium**

- **(A)** Murine intestinal organoids were treated with 0 or 9 Gy and either vehicle or PMX205 1 hour before irradiation. Western blotting was carried with the antibodies indicated. Total Erk1/2 was used as the loading control.
- **(B)** GeneRatios are shown following GO enrichment analysis of murine intestinal organoids treated +/- PMX205 +/- 9 Gy irradiation. Organoids were harvested 48 h post-irradiation. Oxidoreductase activity, acting on … = oxidoreductase activity, acting on single donors with incorporation of molecular oxygen; oxidoreductase activity, acting on single donors with incorporation of molecular oxygen, incorporation of two atoms of oxygen.
- **(C)** GeneRatios are shown following KEGG (Kyoto Encyclopedia of Genes and Genomes) pathway analysis in murine intestinal organoids treated +/- PMX205 +/- 9 Gy irradiation. Organoids were harvested 48 h post-irradiation. Glycosphingolipid biosynthesis – lacto… = Glycosphingolipid biosynthesis - lacto and neolacto series. Glycosaminoglycan biosynthesis – chondroitin… = Glycosaminoglycan biosynthesis - chondroitin sulfate / dermatan sulfate. Glycosphingolipid biosynthesis – globo… = Glycosphingolipid biosynthesis - globo and isoglobo series.

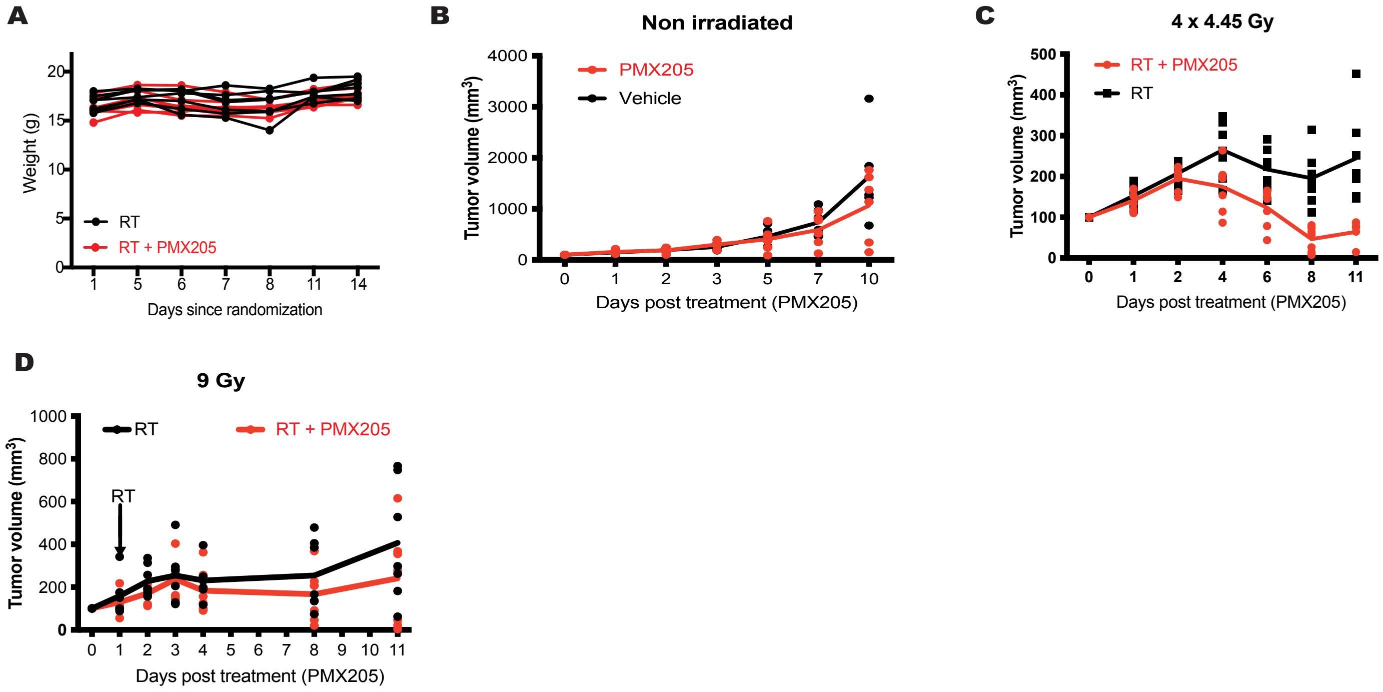
A
Weight (g)
20
15
10
5
0
1 5 6 7 8 11 14
Days since randomization
RT
RT + PMX205
B
Non irradiated
PMX205
Vehicle
Tumor volume (mm$^3$)
4000
3000
2000
1000
0
0 1 2 3 5 7 10
Days post treatment (PMX205)
C
4 x 4.45 Gy
RT + PMX205
RT
Tumor volume (mm$^3$)
500
400
300
200
100
0
0 1 2 4 6 8 11
Days post treatment (PMX205)
D
9 Gy
RT
RT + PMX205
RT
Tumor volume (mm$^3$)
1000
800
600
400
200
0
0 1 2 3 4 5 6 7 8 9 10 11
Days post treatment (PMX205)

**Supplementary Figure 5: C5aR1 inhibition improves tumour radiation response**

- **(A)** Graph shows the weight of mice treated with 9 Gy single dose irradiation and either vehicle or PMX205 treatment for 3 doses flanking the irradiation dose (on day 0, 1 and 2). Individual lines represent individual mice per group.
- **(B)** Tumour growth curves are shown for MC38 subcutaneous tumours treated with either vehicle or PMX205 treatment for 3 doses (on day 0, 1 and 2).
- **(C)** Tumour growth curves are shown for MC38 subcutaneous tumours treated with 3 x 4.45 Gy single dose (equivalent to 9 Gy assuming an $\alpha/\beta$ ratio of 5.06)(Suwinski et al., 2007) and either vehicle or PMX205 treatment for 3 doses flanking the irradiation dose (on day 0, 1 and 2). * = $p<0.05$ , ** = $p<0.01$, *** = $p<0.001$, comparing vehicle and PMX205 treated mice at days 6, 8 and 11 respectively. **** = $p<0.0001$ comparing day 0 to day 11 by 2-way ANOVA with Dunnett's comparison test. Individual points represent individual mice per group.
- **(D)** Tumour growth curves are shown for MC38 subcutaneous tumours treated with 9 Gy single dose irradiation and either vehicle or PMX205 treatment for 3 doses flanking the irradiation dose (on day 0, 1 and 2). **** = $p<0.0001$ comparing vehicle and PMX205 treated mice at day 11. * = $p<0.05$ comparing day 0 to day 11 by 2-way ANOVA with Dunnett's comparison test. Individual points represent individual mice per group.

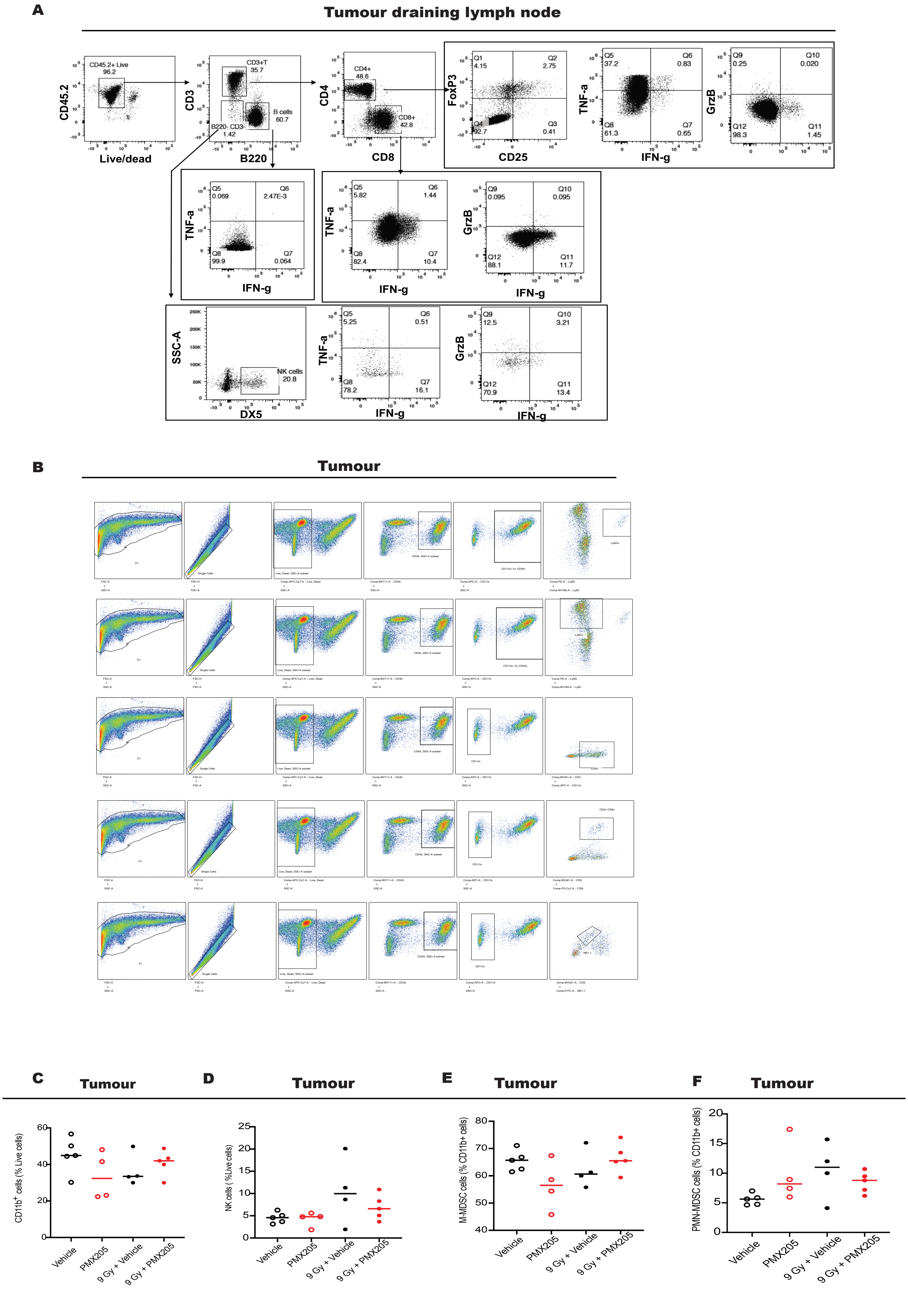
A
Tumour draining lymph node
CD45.2+ Live
96.2
CD45.2
Live/dead
CD3+T
35.7
B cells
60.7
B220- CD3-
1.42
CD3
B220
CD4+
48.6
CD8+
42.8
CD4
CD8
Q1
4.15
Q2
2.75
Q3
0.41
Q4
92.7
FoxP3
CD25
Q5
37.2
Q6
0.83
Q7
0.65
Q8
61.3
TNF-a
IFN-g
Q9
0.25
Q10
0.020
Q11
1.45
Q12
98.3
GrzB
IFN-g
Q5
0.069
Q6
2.47E-3
Q7
0.064
Q8
99.9
TNF-a
IFN-g
Q5
5.82
Q6
1.44
Q7
10.4
Q8
82.4
TNF-a
IFN-g
Q9
0.095
Q10
0.095
Q11
11.7
Q12
88.1
GrzB
IFN-g
SSC-A
DX5
NK cells
20.8
Q5
5.25
Q6
0.51
Q7
16.1
Q8
78.2
TNF-a
IFN-g
Q9
12.5
Q10
3.21
Q11
13.4
Q12
70.9
GrzB
IFN-g
B
Tumour
C
Tumour
CD11b+ cells (% Live cells)
Vehicle
PMX205
9 Gy + Vehicle
9 Gy + PMX205
D
Tumour
NK cells ( %Live cells)
Vehicle
PMX205
9 Gy + Vehicle
9 Gy + PMX205
E
Tumour
M-MDSC cells (% CD11b+ cells)
Vehicle
PMX205
9 Gy + Vehicle
9 Gy + PMX205
F
Tumour
PMN-MDSC cells (% CD11b+ cells)
Vehicle
PMX205
9 Gy + Vehicle
9 Gy + PMX205

**Supplementary Figure 6: Targeting C5aR1 does not increase the % of CD8+ T-cells in the tumour following irradiation**

- **(A)** Gating strategy for the analysis shown in Figures 6B-L.
- **(B)** Gating strategy for the analysis shown in Figures 6M and N (and Supplementary Figures 6C-F).
- **(C)** Graph shows CD11b+ cells (as a % of live cells) in tumours of mice receiving 0 or 9 Gy and either vehicle or PMX205 treatment following the same dosing scheme as shown in Figure 6A. Tumours were harvested 7 days after irradiation with either 0 or 9 Gy. Individual points represent individual mice per group.
- **(D)** Graph shows NK cells (as a % of live cells) in tumours of mice receiving 0 or 9 Gy and either vehicle or PMX205 treatment following the same dosing scheme as shown in Figure 6A. Tumours were harvested 7 days after irradiation with either 0 or 9 Gy. Individual points represent individual mice per group.
- **(E)** Graph shows M-MDSC-cells (as a % of live cells) in tumours of mice receiving 0 or 9 Gy and either vehicle or PMX205 treatment following the same dosing scheme as shown in Figure 6A. Tumours were harvested 7 days after irradiation with either 0 or 9 Gy. Individual points represent individual mice per group.
- **(F)** Graph shows PMN-MDSC cells (as a % of live cells) in tumours of mice receiving 0 or 9 Gy and either vehicle or PMX205 treatment following the same dosing scheme as shown in Figure 6A. Tumours were harvested 7 days after irradiation with either 0 or 9 Gy. Individual points represent individual mice per group.

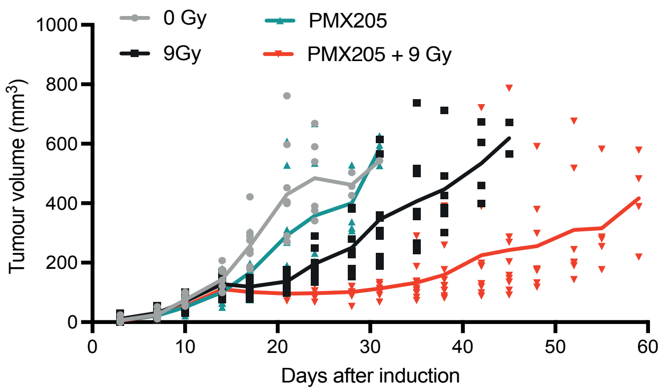

0 Gy
9Gy
PMX205
PMX205 + 9 Gy
Tumour volume (mm3)
1000
800
600
400
200
0
0
10
20
30
40
50
60
Days after induction

**Supplementary Figure 7: C5aR1 inhibition can improve radiotherapy in tumours with an immunosuppressive microenvironment**

Tumour growth curves for AKPT organoids grown subcutaneously and treated with either 0 or 9 Gy and either vehicle or PMX205 flanking irradiation. Individual points = individual mice per group. **** = $p<0.0001$ by 2-way ANOVA with Dunnett's comparison test.

### Supplementary Tables

**Supplementary Table 1:** List of GSEA significant pathways used to plot the data shown in Figure 1E; with corresponding columns denoting which ones are annotated as Complement and Immune System pathways.

**Supplementary Table 2:** Raw data used in the generation of the heatmap shown in Figure 2A, see also methods section.

**Supplementary Table 3:** Pathways differentially expressed by RPPA following treatment with PMX205.

**Supplementary Table 4:** Genes differentially expressed by RPPA following treatment with PMX205.
